# Rapid evolution of pre-zygotic reproductive barriers in allopatric populations

**DOI:** 10.1101/2023.03.18.533249

**Authors:** Anjali Mahilkar, Prachitha Nagendra, Pavithra Venkataraman, Saniya Deshmukh, Supreet Saini

## Abstract

Adaptive divergence leading to speciation is the major evolutionary process generating diversity in life forms. The most commonly observed form of speciation is allopatric speciation which requires that gene flow be prevented between populations by physical or temporal barriers, as they adapt to their respective environments. Eventually, these adaptive responses drive the populations far apart in the genotypic space such that individuals from the two populations become reproductively isolated. A widely accepted theory is that speciation simply occurs as a by-product of adaptive response of the populations^1,2^. Several ecological and laboratory examples of allopatric speciation exist^3–6^. However, we know little about the nature (pre- or post-zygotic) of barriers that arise first in this process. Understanding the first barriers that arise between populations is key, as populations diverge towards becoming distinct species. In recent years, fungi been used as model organisms to answer questions related to evolution of reproductive isolation^3,7–9^. Here we show rapid evolution of pre-zygotic barriers between allopatric yeast populations. We further demonstrate that these pre-zygotic barriers arise due to altered mating kinetics of the evolved population. Moreover, our non-adaptive evolution experiments with yeast under limited selection pressure also show rapid emergence of reproductive isolation. Overall, our results show that evolution of pre-zygotic reproductive barriers can occur as result of natural selection or drift. These barriers result because of altered mating kinetics or mate preference.

**One sentence summary:** Pre-zygotic barriers to gene flow can arise due to adaptation or drift.

## Introduction

How diversity of life forms arise on Earth is an open question. Although, Charles Darwin’s *Origin of Species* explained natural selection as a mechanism for populations to adapt to prevailing environmental conditions, we do not yet understand the fundamental evolutionary forces leading to reproductive isolation. According to one idea, reproductive isolation, leading to speciation, is thought to evolve incidentally as a by-product of adaptation of populations to diverging selection pressures in allopatry^10,11^. Most of the evidence supporting this hypothesis of speciation being a fallout of adaptive response of the population to divergent selection come from study of recently diverged species in ecological environments.

The barriers to gene flow leading to speciation could be pre-zygotic or post-zygotic^12^. However, little is known about the relative importance of the mechanisms and forms of reproductive barriers by which speciation occur. The central challenge that remains in understanding speciation is understanding the nature of and order in which these barriers arise between two populations, eventually leading to speciation^13^. Efforts in this direction come from analysis of closely related species or via laboratory experiments. Populations in allopatry, in the absence of reinforcement or assortative mating, are thought to more readily establish post-zygotic barriers. On the other hand, it is thought that pre-zygotic barriers are more relevant for populations in sympatry. Overall, conclusions regarding the relative roles of pre- and post-zygotic barriers in initiating speciation process are contentious^13–23^.

Experimental evolution, to understand how barriers to gene flow arise, demonstrates that adaptation to ecological niches leads to evolution of reproductive barriers in a relatively short time^24,25^, and can result due to pre-zygotic (pre- ^24^ or post-mating ^26^), or post-zygotic ^27^ barriers. While, these experiments have largely been performed with populations with standing genetic variation ^7,25,26,28–30^, more recently, populations of isogenic yeast has been used a model system to understand evolution of reproductive isolation^3,4,31^. However, the relative contribution of pre- and post-mating barriers in the early steps of speciation remains uncharacterized. In some studies, the experimental design preludes asking this question.

In this work, we ask the following question: which barriers arise first between populations in allopatry, evolving under selection and drift? To answer this question, we perform adaptive and non-adaptive evolutionary experiment with the yeast *S. cerevisiae*. Haploid isogenic yeast populations of both mating types were evolved in limiting glucose and galactose amounts for 600 generations (**Fig. S1**). To study evolution in the absence of selection, haploids were also evolved in a mutation accumulation experiment for 70 transfers (~1500 generations). We demonstrate that metabolic specialization in the adaptive evolution experiment, and the action of drift alone in the non-adaptive experiment leads to rapid evolution of pre-zygotic barriers, in the form of reduced mating efficiency as a result of altered mating kinetics. We demonstrate that, in specific environmental contexts, acquisition of a few SNPs only can establish significant barriers to mating. Overall, we demonstrate that evolution of reproductive barriers can start with pre-zygotic barriers among populations in allopatry.

## Results

### Adaptive response in glucose and galactose

During allopatric evolution, an increase in fitness was observed due to a decrease in the lag phase duration, and an increase in the growth rate (**Fig. 1A and 1B**). The adaptive path of the galactose-evolved lines follows a distinct phenotypic path. In the first 200 generations, the evolved lines exhibit a reduction in the lag phase duration and an increase in the growth rate. In the next 200 generations (200 to 400 generations), the evolved lines exhibit a statistically insignificant increase in both, the growth rate and the lag phase duration. In the last 200 generations of our experiment (400 to 600 generations), a decrease in the lag phase duration was observed. However, this decrease in the lag phase duration was observed at a fitness cost of a decrease in the growth rate. As compared to the galactose-evolved lines, the glucose-evolved haploid lines exhibit a distinctly different pattern. The growth characteristics of the evolved lines when studied in the other environment (i.e., galactose-evolved lines in glucose) is as shown in **Fig. 1C and 1D**.

**Figure 1.**
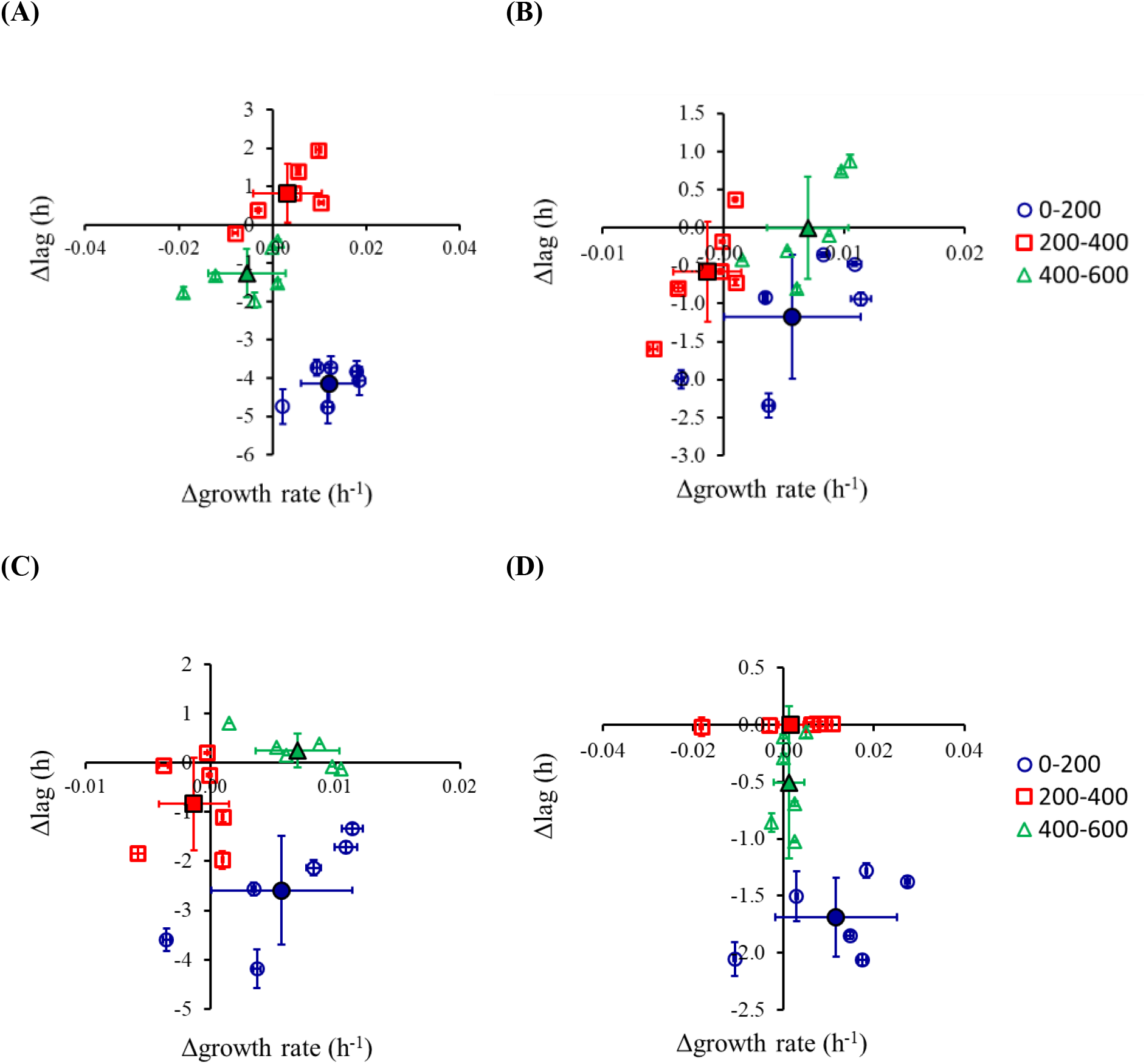
**Adaptive response of haploid yeast during evolution in low galactose (A) and glucose (B).** Changes in growth rate and lag phase duration every 200 generations are shown. **Adaptive response of haploid yeast, in terms of changes in growth rate and lag phase duration, for (C) galactose-evolved lines in glucose, and (D) glucose-evolved lines in galactose.** Open symbols are independent lines. Closed symbols are average of the six lines. Growth experiments were performed three independent times. The average and standard deviation is shown.

### Pre-zygotic barriers to mating evolve rapidly

Reproductive barriers between populations can be quantified via studying three processes in yeast: change in mating efficiency, mitotic growth of the hybrid, and meiotic efficiency of the hybrid. The pairwise mating efficiencies between all evolved haploid lines exhibit a marked decrease, as compared to the ancestor (**Fig. 2 and S2**). This trend is observed for both, ecological speciation as well as for mutation-order speciation. The decrease in the mating efficiency is most rapid between the glucose-evolved lines.

**Figure 2.**
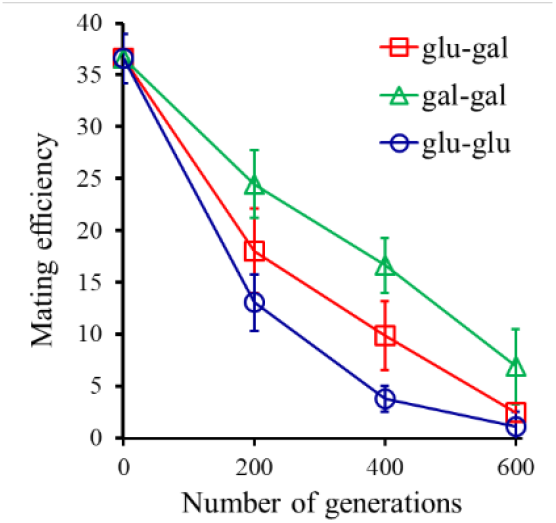
Mating efficiency between the evolved lines evolved in glucose- or galactose environments. The data represents the average of nine possible pairings for each condition, and time point. All experiments were performed three independent times. The average values and the standard deviation are shown in the Figure.

The decrease in mating efficiency between evolved haploids could possibly be due to one of two reasons. First, the mitotically-evolving lines diverged genetically and as a result, despite retaining intact mating pathways, no longer mate with each other with the same efficiency. Second, since these lines were evolved without selection pressure to retain the genes necessary for mating, these lines acquired mutations, which rendered them incapable of mating. To distinguish which one of these two possibilities play out in our evolved populations, we switched the mating type of each of the haploid lines at 600 generations. For example, the mating type of galactose evolved **a** was switched to **α**. Similar mating type switch was done for all haploid lines. We then quantified the mating efficiency of each evolved haploid with self (carrying the opposite mating type). Each of the evolved haploids mate with their mating-type switched counterpart with considerably higher efficiency, as compared with the average mating efficiency of the evolved haploid with all other evolved haploid lines (**Fig. 3**). These results clearly indicate that the evolved haploids retain intact mating pathways. Lines glu1a and glu2a were left out from this analysis, as these lines had undergone an autodiploidization during the course of the experiment (**Fig. S3**) ^32,33^.

**Figure 3.**
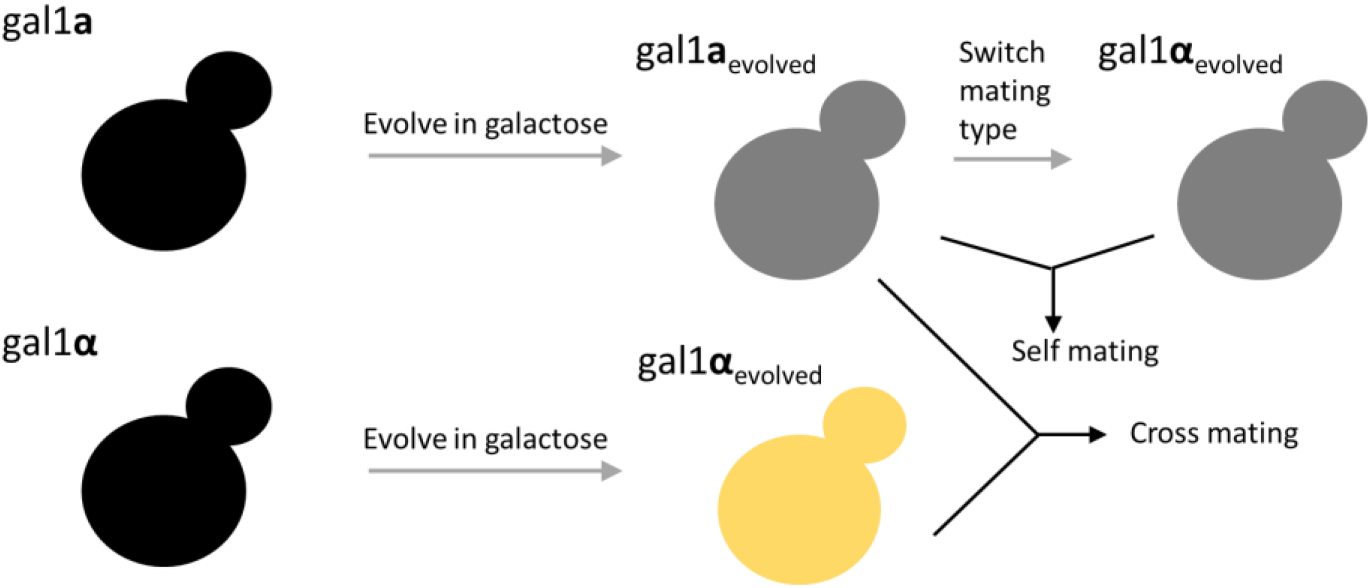
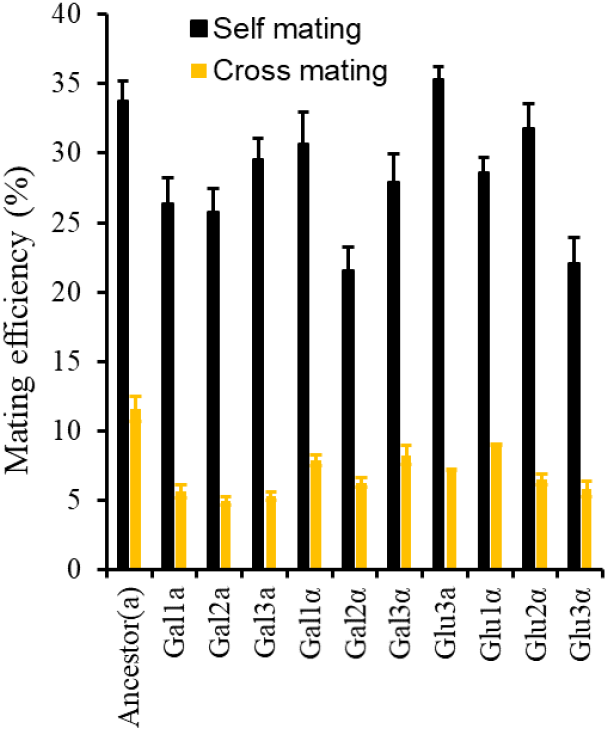
**(Top)** Schematic of cross- and self-mating of an evolved haploid. **(Bottom)** Mating efficiency of the evolved haploids with their mating switched (black bars). Mating efficiency of an evolved haploid with itself (mating type switched) is greater than mating with other haploids of the opposite mating type (yellow bars). Cross mating efficiency represents the average of the mating efficiency with all other six evolved haploids of the opposite mating type and the ancestor. All experiments were performed three independent times. The average and the standard deviation is presented in the figure.

From the 12 evolved haploid lines and the ancestor **a** and **α**, 49 hybrids could be possibly generated. Of all these possibilities, we were only able to create 42 (**Fig. S4**). Seven hybrids, all from the glu1a and glu2a lines, could not be created. Thereafter, we characterized the growth kinetics of the 42 hybrid lines in glucose and galactose environments.

### Nature of epistasis between beneficial mutations in glucose and galactose adapted lines

The hybrids were separated into five groups. Those resulting from mating between (i) two haploids evolved in glucose (glu-glu), (ii) haploids evolved in galactose (gal-gal), (iii) a glucose- and a galactose-evolved haploid (glu-gal), (iv) a glucose-evolved haploid with an ancestral haploid (glu-anc), and (v) a galactose-evolved haploid with an ancestral haploid (gal-anc). The performance of all five groups of hybrids was compared with that of the ancestral diploid.

In glucose, the hybrids in the glu-glu, glu-anc, and the glu-gal group, all exhibited a decrease in the growth rate, compared to the ancestral diploid (**Fig. 4A, 4B, and S5; Tables S1 and S2**). This growth defect is most severe in the glu-glu diploids. Thus, bringing together adaptive mutations in a single genome has a detrimental effect on cellular fitness. This effect has previously been seen during growth in glucose, however, the precise mechanistic details remain unknown^34^. The fitness effect of epistatic interactions in yeast is known^35,36^, and its possible role as a mechanism leading to speciation has been suggested recently^37^. On the other hand, the hybrids exhibit a qualitatively different pattern, when grown in galactose. The gal-gal group of hybrids exhibits the greatest growth rate, and among the shortest lag phase duration. These results suggest that epistasis between beneficial mutations in glucose is qualitatively different from that in galactose. In addition, a small number of hybrids exhibit a statistically significant decrease in the meiotic efficiency, as compared to the ancestor (**Fig. S6**).

**Figure 4.**
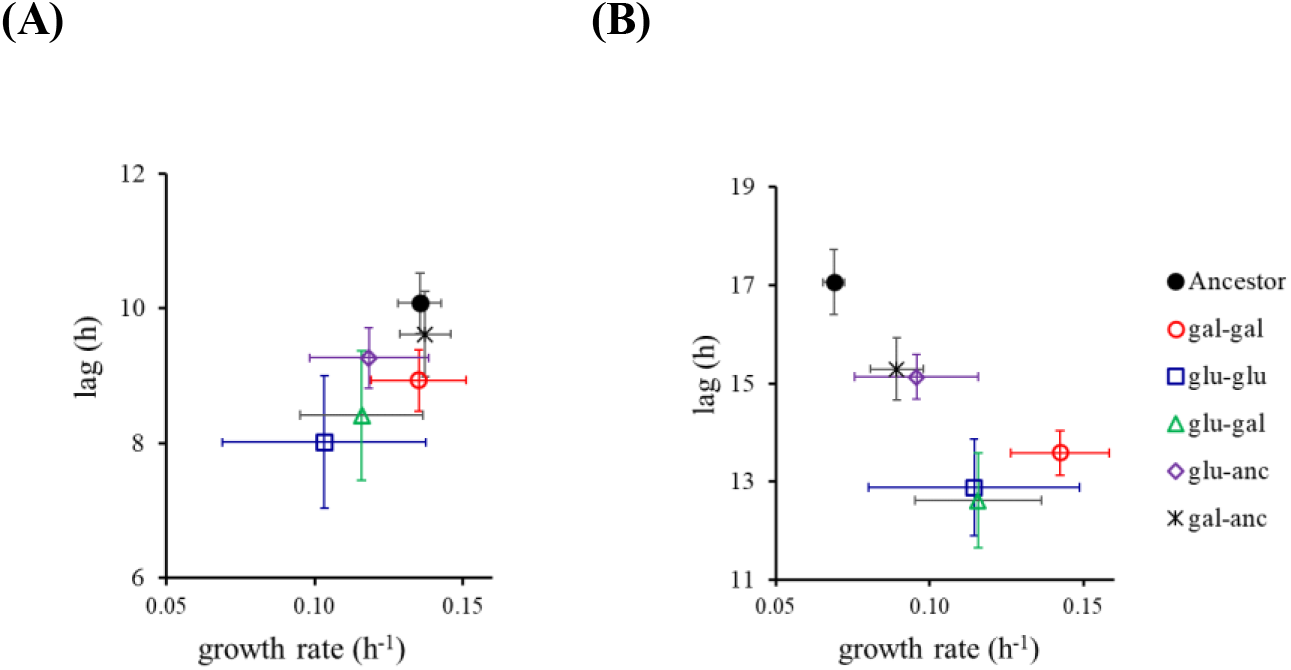
**Mitotic performance of the hybrids in glucose (A) and galactose (B).** The hybrids are separated in five groups, depending on the ancestral haploids. Average and standard deviation of three independent experiments is shown. See Tables S1 and S2 for statistics between the five groups, and Fig. S5 for individual data points.

Overall, our data suggests that, in allopatry, pre-zygotic barriers arise significantly faster as compared to post-zygotic barriers. In the framework of our evolution experiment, adaptation has two components (a) decrease in lag phase duration and (b) increase in growth rate. No correlation exists between these values of components of fitness and that of pre- or post-zygotic barriers to gene flow.

### Genome sequencing reveals signatures of convergent evolution

To identify the genetic basis of adaptation and reproductive isolation between the evolved haploid lines, we sequenced the genomes of the 12 haploid lines, after adaptation for 600 generations (**Fig. S7, Table S3, Annexure I**), and compared with the ancestral sequence. The sequencing results show evidence of convergent evolution. Two glucose-evolved lines have mutations in MNS1, an ER membrane protein responsible for α-mannosidase activity (a stop codon and a frameshift mutation). MNS1, mutants exhibit a significantly longer life span in yeast^38^, and in other organisms^39^. Two galactose-evolved lines have mutations in MNL1 (a stop codon and a non-synonymous mutations), another Mannosidase-like protein, which works via formation of a complex with the protein disulfide isomerase, PDI1^40,41^. Two lines (one glucose- and one galactose-evolved) have point mutations (both synonymous) in the gene PRM7. PRM7p is induced several-fold in the presence of pheromone, and is thought to be involved in cell-cell communication^42,43^.

### Evolved lines exhibit altered kinetics of mating

While these similarities in genic targets exist, considerable difference in the mutational targets are also present between the different evolved lines (**Table S3**). These results lead us to hypothesize that altered mating kinetics in the evolved lines lead to establishment of first reproductive barriers. To test this possibility, we perform mating experiments where kinetics of mating between evolved haploids **a** and their counterparts **α** (obtained by their mating switched). As shown in **Fig. 5**, the evolved haploids, when mated with their opposite mating type (obtained by switching mating type), exhibit (a) no mating defect, and (b) altered mating kinetics. This observation lends further strength to the argument that reproductive barriers between the haploid evolved lines (and the ancestor) are largely a result of altered program of the mating kinetics for a cell.

**Figure 5.**
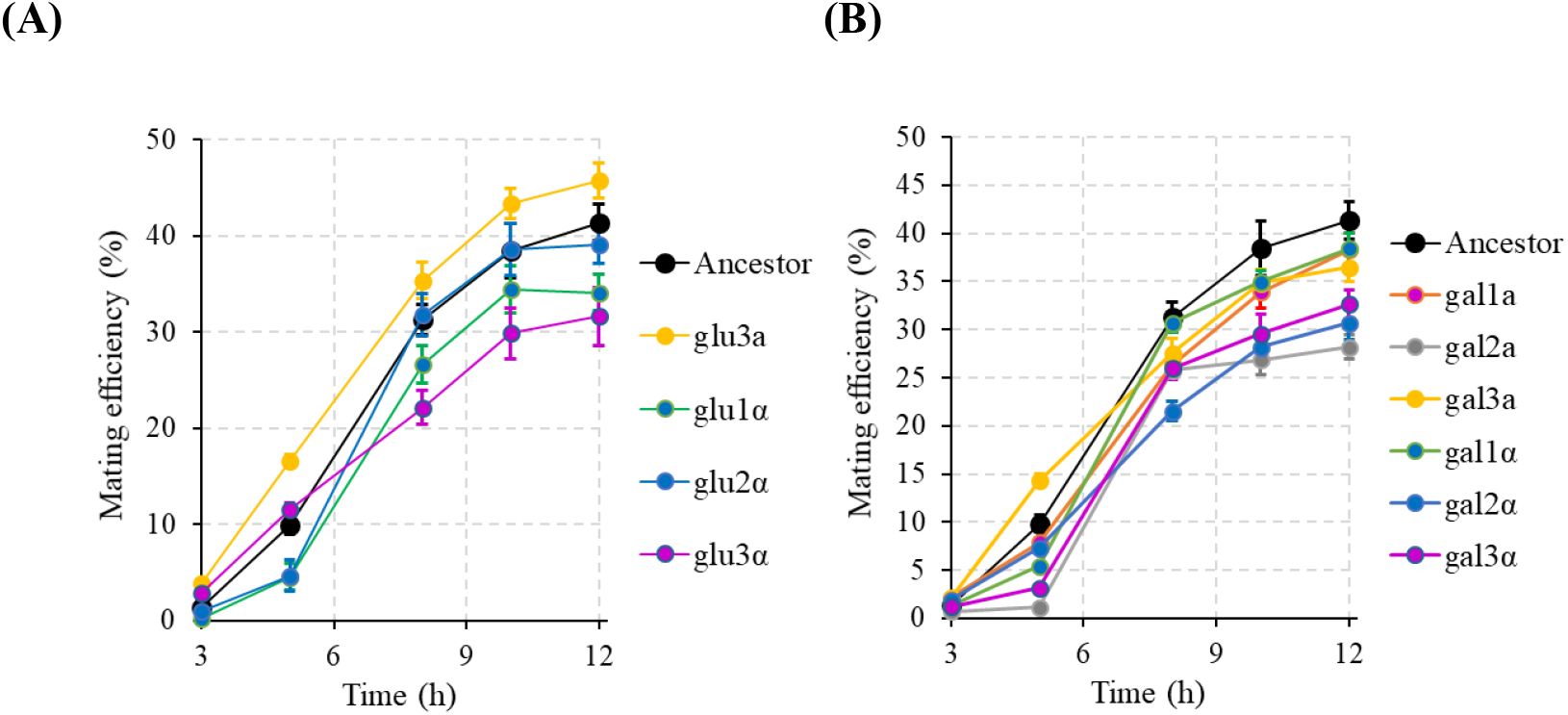
**Kinetics of mating efficiency between (A) glucose-evolved haploids and the switched mating partner; (B) galactose-evolved haploids and the switched mating partner.** Ancestor mating efficiency data is between the ancestral **a** and ancestral **α**. All experiments were performed in triplicate. Average values and the standard deviation are reported. In (A), other than glu2α, all profiles are statistically different from that of the ancestor (p-value < 0.05). In (B), all profiles, other than gal1α, are statistically different from that of the ancestor (p-value < 0.05).

### Pre-zygotic barriers arise first in allopatric populations evolved under drift

The null model of speciation suggests that speciation results as a by-product of an adaptive process. Our data above also suggests this. However, this hypothesis has never been explicitly tested in an experimental context. In order to do this, we evolved 22 haploid lines **a** and **α** each in a mutation accumulation (MA) experiment. The 44 lines (a1-a22 and α1-α22) were propagated for 70 transfers (~1500 generations). Since a severe bottleneck is imposed at every transfer, the role of selection is largely absent, and drift dictates evolutionary trajectory. MA experiments with microbial systems demonstrate that, under constant propagation under these conditions leads to a reduction in the fitness of MA lines ^44^. After 70 transfers, the mean fitness of the evolved lines is ~0.98 times that of the ancestor (**Fig. S8**).

Our results show that after 70 transfers, a few lines exhibit a statistically significant decrease in the mating efficiency with the ancestor haploid. This decrease in mating efficiency has taken place in the absence of adaptation, and is present in varying degrees (ranging from a decrease of ~40% to an increase of ~4%), among the 44 lines evolved. There is no correlation between the adaptive response of the MA lines, and the extent of change in the mating efficiency with the ancestral haploid. These results clearly demonstrate that reproductive isolation can, in addition to resulting from adaptation, also result as a by-product of drift (**Fig. 6**).

**Figure 6.**
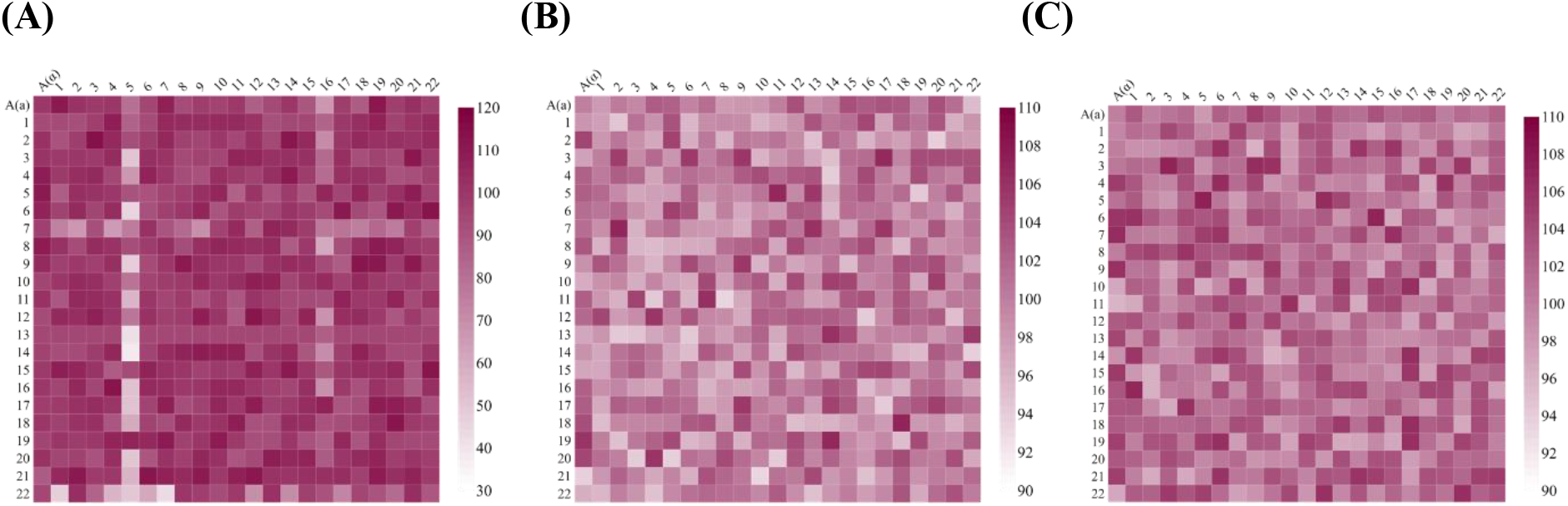
Reproductive isolation emerges most rapidly via pre-mating barriers for haploids evolving under drift. **(A) Mating efficiency of MA lines of α (x-axis) and a (y-axis). (B)** Mitotic growth rate of the hybrids, as compared to the ancestor. **(C)** Meiotic efficiency of the hybrids, as compared to the ancestor. For (A), (B), and (C), the ancestral mating efficiency, mitotic growth rate, and meiotic efficiency respectively is normalized to 100. Numbers “1” to “22” on the x- and the y-axis represent the 22 evolved haploid lines. “A” refers to the ancestor. All experiments were performed three independent times. The average of the three is reported here. The standard deviation for each data point is less than 10% of the data value. The ancestral mating efficiency (A), mitotic growth rate (B), and meiotic efficiency (C) is normalized to 100 in each of the three plots. Also see Fig. S9 for the three plots with identical range for the heat plot.

## Discussion

Speciation is the fundamental process that generates diversity of life forms on Earth. The most widely accepted explanation for speciation is that it occurs as a fallout of adaptation to diverging selection. Evolution of reproductive barriers among individuals in a population is a classic long-standing interest among biologists trying to understand the process of speciation. However, which barriers arise first as groups begin to diverge in allopatry?

Our data suggests that upon starting with isogenic populations, adaptation in allopatry leads to rapid evolution of pre-zygotic barriers in yeast. In addition, drift alone can also lead to establishment of pre-zygotic barriers. Mating kinetics in yeast are intricately controlled^45^. In fact, not only adaptation, but metabolic state of isogenic cells controls the effectiveness and selection of a mating partner^8^. In context of adaptation to galactose, different ecological contexts lead to non-random segregation of GAL alleles among environmental isolates^46^. Additionally, alleles of galactose utilization regulators GAL4 and GAL80 have been isolated in laboratory^47^, leading to altered growth and fitness in different environments ^48^. In specific environmental contexts, these allelic distributions in an otherwise isogenic background is sufficient to establish a pre-zygotic barrier (**Fig. S10**).

Reproductive barriers could arise in several ways. Pre-zygotic barriers have been known to arise rapidly, in response to divergent selection^49,50^, although a few negative examples are known too^51,52^. Overall, however, pre-zygotic isolation, as a by-product of adaptation, is observed quite frequently^25^. Post-zygotic barriers can be (a) universal^53–55^ among closely-related species but are likely rare in the initial stages of speciation and hence, have not been observed in laboratory experiments; or (b) environment-dependent in nature. Regarding (b), while negative epistasis widely observed among beneficial mutations^35,36,56^, sign-epistasis is relatively infrequent. Antagonism due to a beneficial mutation is also strongly dependent on the choice of the two environments^57–59^. These results suggest that while post-zygotic barriers could arise due to multiple mechanisms and have been predicted to arise in theory too^37^, it is likely that pre-zygotic barriers arise rapidly in allopatry due to behavioral changes.

## Materials and Methods

### Methods

#### Strain construction

*Saccharomyces cerevisiae* strain used in this work is a derivative of ScPJB644 (*MATa MEL1 ade1 ile trp1-HIII ura3-52)^60^* with auxotrophic markers inserted near MAT locus to identify the mating type of the haploids. To prepare **a** and ***α*** ancestral population, ScPJB644 diploid was sporulated and dissected to obtain isogenic **a** and ***α*** haploids from a single ascospore. Two different auxotrophic markers, TRP1 and URA3, were inserted at the same site in ARS314 on chromosome III near the MAT locus on each **a** and ***α*** type haploids respectively. They are located between genes PHO87 and BUD5, but not disrupting either gene (**Fig. S12**), as described earlier^61^.

URA3 was amplified from the plasmid p426GPD^60^ using the primer set pSc011 and pSc012 (all primers listed in **Table S4**). TRP1 was amplified from the plasmid p424TEF^62^ using the primer set pSc014 and pSc015. Both URA3 and TRP1 fragments were further to two sequential rounds of PCR to increase the length of flanking ends for efficient recombination, using the primer sets pSc018 and pSc019, and pSc020 and pSc021. Purified PCR products were transformed into **a** and ***α*** type haploids by electroporation, using Eppendorf eporator. The two haploids thus obtained are referred to as ScAM04 (*MATa URA3*) and ScAM05 (*MATa TRP1*). The ancestral diploid was obtained by mating the two and the resulting strain is referred to as ScAM06 (*MATa TRP1/MAT α URA3*). All strains were grown in YPD (0.5% Yeast Extract, 1% Peptone and 2% Dextrose (w/v)) at 30°C and shaking at 250 rpm in 25×100 mm test-tubes unless specified otherwise. The freezer stocks of all strains are stored at −80°C in 25% (v/v) glycerol.

The GAL4c GAL80s-1 strains used in this work are described in^47,63^.

#### Evolution experiment

The ancestral population for the evolution experiment were started from the freezer stocks streaked on YPD plates. Single colonies of each ScAM04 and ScAM05 were inoculated in 5ml YPD for 36 hours at 30°C and shaking at 250 rpm. These cultures were used as inoculum for the evolution experiment. Three replicate lines of each haploid strains were started in 0.2% (w/v) glucose or galactose in standard Synthetic Complete Medium (SCM: 0.671% YNB with nitrogen base and 0.05% complete amino acid mixture). Thus, in total, we maintain 12 populations, three of each kind.

Adaptive evolution experiments were performed by serial dilutions every 24 hours in SCM with appropriate carbon source. Every 24 hours, growing cultures were diluted 1:100 in fresh SCM medium with appropriate carbon source yielding ~6-7 generations every 24 hours. Intermediate generations were cryopreserved in 25% (v/v) glycerol every 200 generations.

#### Mutation accumulation (MA) experiment

ScAM04 was spread on YPD plates for single colonies. Prior to spreading, a small area was identified and marked on the plate. The colony inside (or closest to) the marked area, after 48 hours of growth at 30 °C was suspended in 2 ml PBS buffer. A fraction of this volume was spread on a fresh YPD plate for single colonies, and the process is repeated for 70 transfers. A total of 22 independent lines of ScAM04 were maintained in this experiment. Similarly, 22 independent lines for ScAM05 were propagated for the MA experiment.

#### Fitness measurements

Evolved lines and ancestor (a or *α* haploids) from freezer stocks were revived in 5ml YPD and incubated for 48 hours then transferred 1:100 to glycerol-lactate medium (gly-lac: 3% Glycerol, 2% potassium lactate (pH 5.6), 0.671% YNB with nitrogen base and 0.05% complete amino acid mixture) and incubated for 48 hours at 30° C and shaking at 250 rpm. 2ml SCM with appropriate carbon source (0.2% glucose/galactose) were inoculated for a final OD_600_ of 0.01. 150μl of the cultures were then transferred to a clear flat-bottom 96-well plate (Costar) in triplicates and incubated at 30°C in an automated microplate reader (Tecan Infinite M200 Pro) until they reached stationery phase. A gas permeable Breathe-Easy (Sigma-Aldrich) sealing membrane was used to seal the 96-well plates. OD600 measurements were taken every 1 hour with 15 minutes of orbital shaking at 5 mm amplitude before the readings. Growth rates were calculated by plotting log (OD) from the exponential phase of growth against time. The slope of the straight-line fit was determined as the growth rate of the strain. The point of intersection of this straight line, with the straight line with the equation log (initial OD) was taken as the duration of the lag phase.

#### Mating efficiency

Ancestor and evolved haploid lines were revived in 5ml YPD cultures from freezer stocks and grown till saturation at 30°C. The haploids of opposite mating type were spot-plated on a 1cm^2^ area leaving 1 cm space between the two and incubated at 30 °C for 24 hours. After incubation, cells from either side of the square were scraped and resuspended in sterile water and measured OD600 to determine cell density. Equal number of both the haploids were mixed in a 1.5 ml sterile microcentrifuge tubes and 5 ul of the mix were spot-plated in center 1 cm^2^ marked area and allowed mating for 7 hours at 30 °C. After 7 hours, cells from this area were scraped and plated on YPD plates for single colonies and incubated at 30 °C for 24 hours (**Fig. S13**). The diploids were counted by replica-plating on Uracil and Tryptophan double dropout synthetic medium agar plates. A minimum of 500 colonies were transferred for each mating efficiency to the double dropout plate, for quantification of the mating efficiency. Mating efficiency was determined by the formula,

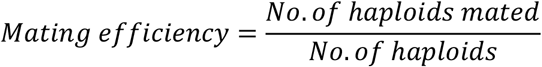

#### Sporulation efficiency

The hybrids or homozygous diploids, along with the ancestral diploid were revived by streaking on YPD plates from freezer stocks. Single colonies from each line were patched on to freshly prepared pre-sporulation GNA plates (GNA medium: 5% D-glucose, 3% nutrient broth, 1% yeast extract, and 2% agar) and incubated at 30°C for 24 hours. Re-patched again on GNA plates for another 24 hours. Small lumps of cells were then patched onto sporulation medium plates containing 1% Potassium acetate and 2% Agar and incubated at 25°C for 5 days and 30° C for 3 days. Sporulated/ un-sporulated cells were directly counted under ×400 magnification using microscope to determine the sporulation efficiency (a minimum of 500 cells). The following formula was used to determine the sporulation efficiency.

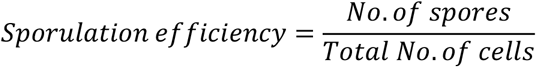

#### Self-mating efficiency of the evolved lines

Mating type **a** evolved haploid lines after 600 generations were transformed with plasmid carrying the *HO* gene to obtain diploids of the evolved lines by selecting on URA dropout plates. The diploids were then sporulated as described previously. Tetrad dissection was performed via standard zymolase treatment and manually dissection under the microscope. Haploids of both mating types were obtained from a single ascus and identified using colony PCR and primers described in^64^. We then performed mating assay as described previously and used colony PCR of the *MAT* locus to determine the number of haploids and diploids to determine the mating efficiency. At least 150 colonies were analysed using PCR to quantify the mating efficiency between two haploids in a single experiment. Self-mating efficiency reported here is an average of three independent self-mating experiments for all pairs analysed.

#### Mutation accumulation experiment

Cells from a single founder colony of **a** mating type were taken, and spread of YPD plates (containing 1% glucose). Prior to spreading on the YPD plate a random area was marked on the plate. A colony which arose closest to the marked area was taken for propagation in the experiment. The same process was repeated for 22 lines of the **a** haploid independently. Seventy transfers were completed for each of the 22 lines. Each colony has about 10^8^ cells, and therefore, each transfer corresponds to about 25 generations.

#### FACS experiment

A 5ml YPD culture was started from a single colony on a YPD plate. Saturated cultures were sub-cultured 1:100. When the cultures reach an OD of 1.00, a volume of 1.5 ml of the culture in exponential phase was spun down (2-3 x 10^6^ cells/ml) and resuspended in 100 ul autoclaved water. A volume of 1 ml of 70% ethanol was added along the sides of the vial while vortexing. The cells were then incubated at room temperature for an hour and then kept at 4°C overnight. The cells were then washed with 500ul of RNase buffer (0.2M Tris-Cl with 20mm EDTA, pH 8.0) and resuspended in 100ul of the same buffer. RNase A was added to a final concentration of 1 ug/ul. The cells were then incubated at 27°C for 4 hours, and then washed with 1 ml PBS (centrifuge at maximum speed for 45 sec) and resuspended in 950 ul of the same buffer. 50ul of 1 mg/ml propidium iodide was added to a final concentration of 50ug/ml. This was incubated at room temperature for 30 min. A volume of 500 ul of the cell suspension was gently vortexed and sonicated before analysis through flow cytometer (Becton Dickinson, FACS Aria SORP).

#### Statistical Tests

To compare different sets of data, two-tailed independent samples t-tests were performed. The p-values corresponding to 4 degrees of freedom were obtained to identify difference in the data due to chance.

#### Whole Genome Sequencing and Variant Calling

##### Sample preparation and sequencing

Genomic DNA of ancestral haploid and evolved lines were isolated following standard zymolase based protocol using the kit from *FAVORGEN Biotech Corporation*. DNA concentrations and quality were measured using a Nano-spectrophotometer from eppendorf and by gel electrophoresis. Samples were sent for paired-end sequencing on an Illumina HiSeq, with an average read length of 150 bp. Each of the samples had a minimum coverage of 100x. The raw sequencing data is available at https://www.ncbi.nlm.nih.gov/sra/PRJNA767895.

##### Mapping and variant calling

We used cloud-based web interface system Galaxy (https://usegalaxy.eu/) to perform all sequence data analysis. Illumina paired-end reads were uploaded into the server. The quality of the reads was assessed using FastQC (Version 0.72). Raw reads were trimmed with Trimmomatic (Version 0.38.1)^65^ and mapped to S288c genome (version R64) and variant calling was performed using the automated tool Snippy (Version 4.5.5), according to the best practices recommendations for evaluating single nucleotide variant calling^66^. Variants present in the ancestral strain were filtered out manually. Finally, all remaining indels and SNPs were verified using intensive manual curation.

## Acknowledgements

AM, SS thank ICTS Bangalore for support towards attending the IV^th^ Population Genetics and Evolution School, held in 2020. **Funding:** DBT/Wellcome Trust (India Alliance), grant number IA/S/19/2/504632 (SS, PN). Council of Scientific and Industrial Research (CSIR), Government of India, as a Senior Research Fellow (09/087(0873)/2017-EMR-I) (AM). Institute Post Doctoral Fellowship Program, IIT Bombay (SD). **Author Contributions:** Conceptualization: AM, SS; Methodology: AM, PN, PV, SD, SS; Investigation: AM, PN, PV, SD, SS; Funding Acquisition: AM, SD, SS; Project Administration: SS; Supervision: AM, SS; Writing: AM, SS. **Competing interests:** The authors declare that they have no competing interests related to this study. **Data and Materials Availability:** The raw sequencing data is available at https://www.ncbi.nlm.nih.gov/sra/PRJNA767895.

## Notes

### Competing Interest Statement

The authors have declared no competing interest.

## References

1 Schluter, D. Evidence for ecological speciation and its alternative. Science 323, 737–741, doi:10.1126/science.1160006 (2009).

2 Nosil, P. & Flaxman, S. M. Conditions for mutation-order speciation. Proc Biol Sci 278, 399–407, doi:10.1098/rspb.2010.1215 (2011).

3 Anderson, J. B. et al. Determinants of divergent adaptation and Dobzhansky-Muller interaction in experimental yeast populations. Curr Biol 20, 1383–1388, doi:10.1016/j.cub.2010.06.022 (2010).

4 Dettman, J. R., Sirjusingh, C., Kohn, L. M. & Anderson, J. B. Incipient speciation by divergent adaptation and antagonistic epistasis in yeast. Nature 447, 585–588, doi:10.1038/nature05856 (2007).

5 Thacker, C. E. Patterns of divergence in fish species separated by the Isthmus of Panama. BMC Evol Biol 17, 111, doi:10.1186/s12862-017-0957-4 (2017).

6 Knowlton, N., Weigt, L. A., Solorzano, L. A., Mills, D. K. & Bermingham, E. Divergence in proteins, mitochondrial DNA, and reproductive compatibility across the isthmus of Panama. Science 260, 1629–1632, doi:10.1126/science.8503007 (1993).

7 Dettman, J. R., Anderson, J. B. & Kohn, L. M. Divergent adaptation promotes reproductive isolation among experimental populations of the filamentous fungus Neurospora. BMC Evol Biol 8, 35, doi:10.1186/1471-2148-8-35 (2008).

8 Smith, C., Pomiankowski, A. & Greig, D. Size and competitive mating success in the yeast Saccharomyces cerevisiae. Behav Ecol 25, 320–327, doi:10.1093/beheco/art117 (2014).

9 Tusso, S., Nieuwenhuis, B. P. S., Weissensteiner, B., Immler, S. & Wolf, J. B. W. Experimental evolution of adaptive divergence under varying degrees of gene flow. Nat EcolEvol 5, 338–349, doi:10.1038/s41559-020-01363-2 (2021).

10 Gavrilets, S. Perspective: models of speciation: what have we learned in 40 years? Evolution 57, 2197–2215, doi:10.1111/j.0014-3820.2003.tb00233.x (2003).

11 Schluter, D. Ecology and the origin of species. Trends Ecol Evol 16, 372–380, doi:10.1016/s0169-5347(01)02198-x (2001).

12 Bozdag, G. O. & Ono, J. Evolution and molecular bases of reproductive isolation. Curr Opin Genet Dev 76, 101952, doi:10.1016/j.gde.2022.101952 (2022).

13 Orr, J. A. C. a. H. A. Speciation. (Sinauer Associates (Oxford University Press), 2004).

14 Presgraves, D. C. & Meiklejohn, C. D. Hybrid Sterility, Genetic Conflict and Complex Speciation: Lessons From the Drosophila simulans Clade Species. Front Genet 12, 669045, doi:10.3389/fgene.2021.669045 (2021).

15 Coyne, J. A. & Orr, H. A. “Patterns of Speciation in Drosophila” Revisited. Evolution 51, 295–303, doi:10.1111/j.1558-5646.1997.tb02412.x (1997).

16 Grant, P. R. & Grant, B. R. Genetics and the origin of bird species. Proc Natl Acad Sci U S A 94, 7768–7775, doi:10.1073/pnas.94.15.7768 (1997).

17 Coughlan, J. M. & Matute, D. R. The importance of intrinsic postzygotic barriers throughout the speciation process. Philos Trans R Soc Lond B Biol Sci 375, 20190533, doi:10.1098/rstb.2019.0533 (2020).

18 Turissini, D. A., McGirr, J. A., Patel, S. S., David, J. R. & Matute, D. R. The Rate of Evolution of Postmating-Prezygotic Reproductive Isolation in Drosophila. Mol Biol Evol 35, 312–334, doi:10.1093/molbev/msx271 (2018).

19 Christie, K. & Strauss, S. Y. Along the speciation continuum: Quantifying intrinsic and extrinsic isolating barriers across five million years of evolutionary divergence in California jewelflowers. Evolution 72, 1063–1079, doi:10.1111/evo.13477 (2018).

20 Lackey, A. C. & Boughman, J. W. Evolution of reproductive isolation in stickleback fish. Evolution 71, 357–372, doi:10.1111/evo.13114 (2017).

21 Abzhanov, A., Protas, M., Grant, B. R., Grant, P. R. & Tabin, C. J. Bmp4 and morphological variation of beaks in Darwin’s finches. Science 305, 1462–1465, doi:10.1126/science.1098095 (2004).

22 Scopece, G., Musacchio, A., Widmer, A. & Cozzolino, S. Patterns of reproductive isolation in Mediterranean deceptive orchids. Evolution 61, 2623–2642, doi:10.1111/j.1558-5646.2007.00231.x (2007).

23 Jewell, C., Papineau, A. D., Freyre, R. & Moyle, L. C. Patterns of reproductive isolation in Nolana (Chilean bellflower). Evolution 66, 2628–2636, doi:10.1111/j.1558-5646.2012.01607.x (2012).

24 Villa, S. M. et al. Rapid experimental evolution of reproductive isolation from a single natural population. Proc Natl Acad Sci U S A 116, 13440–13445, doi:10.1073/pnas.1901247116 (2019).

25 Rice, W. R. & Hostert, E. E. Laboratory Experiments on Speciation: What Have We Learned in 40 Years? Evolution 47, 1637–1653, doi:10.1111/j.1558-5646.1993.tb01257.x (1993).

26 Turner, E., Jacobson, D. J. & Taylor, J. W. Genetic architecture of a reinforced, postmating, reproductive isolation barrier between Neurospora species indicates evolution via natural selection. PLoS Genet 7, e1002204, doi:10.1371/journal.pgen.1002204 (2011).

27 S, M. G. & Joshi, A. Evolution of reproductive isolation as a by-product of divergent life-history evolution in laboratory populations of Drosophila melanogaster. Ecol Evol 2, 3214–3226, doi:10.1002/ece3.413 (2012).

28 Duffy, S., Burch, C. L. & Turner, P. E. Evolution of host specificity drives reproductive isolation among RNA viruses. Evolution 61, 2614–2622, doi:10.1111/j.1558-5646.2007.00226.x (2007).

29 Mooers, A. O., Rundle, H. D. & Whitlock, M. C. The Effects of Selection and Bottlenecks on Male Mating Success in Peripheral Isolates. Am Nat 153, 437–444, doi:10.1086/303186 (1999).

30 Rundle, H. D., Chenoweth, S. F., Doughty, P. & Blows, M. W. Divergent selection and the evolution of signal traits and mating preferences. PLoS Biol 3, e368, doi:10.1371/journal.pbio.0030368 (2005).

31 Leu, J. Y. & Murray, A. W. Experimental evolution of mating discrimination in budding yeast. Curr Biol 16, 280–286, doi:10.1016/j.cub.2005.12.028 (2006).

32 Tung, S., Bakerlee, C. W., Phillips, A. M., Nguyen Ba, A. N. & Desai, M. M. The genetic basis of differential autodiploidization in evolving yeast populations. G3 (Bethesda) 11, doi:10.1093/g3journal/jkab192 (2021).

33 Gerstein, A. C., Chun, H. J., Grant, A. & Otto, S. P. Genomic convergence toward diploidy in Saccharomyces cerevisiae. PLoS Genet 2, e145, doi:10.1371/journal.pgen.0020145 (2006).

34 Kvitek, D. J. & Sherlock, G. Reciprocal sign epistasis between frequently experimentally evolved adaptive mutations causes a rugged fitness landscape. PLoS Genet 7, e1002056, doi:10.1371/journal.pgen.1002056 (2011).

35 Costanzo, M. et al. A global genetic interaction network maps a wiring diagram of cellular function. Science 353, doi:10.1126/science.aaf1420 (2016).

36 Kuzmin, E. et al. Systematic analysis of complex genetic interactions. Science 360, doi:10.1126/science.aao1729 (2018).

37 Dagilis, A. J., Kirkpatrick, M. & Bolnick, D. I. The evolution of hybrid fitness during speciation. PLoS Genet 15, e1008125, doi:10.1371/journal.pgen.1008125 (2019).

38 Fabrizio, P. et al. Genome-wide screen in Saccharomyces cerevisiae identifies vacuolar protein sorting, autophagy, biosynthetic, and tRNA methylation genes involved in life span regulation. PLoS Genet 6, e1001024, doi:10.1371/journal.pgen.1001024 (2010).

39 Liu, Y. L. et al. Reduced expression of alpha-1,2-mannosidase I extends lifespan in Drosophila melanogaster and Caenorhabditis elegans. Aging Cell 8, 370–379, doi:10.1111/j.1474-9726.2009.00471.x (2009).

40 Pfeiffer, A. et al. A Complex of Htm1 and the Oxidoreductase Pdi1 Accelerates Degradation of Misfolded Glycoproteins. J Biol Chem 291, 12195–12207, doi:10.1074/jbc.M115.703256 (2016).

41 Sakoh-Nakatogawa, M., Nishikawa, S. & Endo, T. Roles of protein-disulfide isomerase-mediated disulfide bond formation of yeast Mnl1p in endoplasmic reticulum-associated degradation. J Biol Chem 284, 11815–11825, doi:10.1074/jbc.M900813200 (2009).

42 Logsdon, B. A. & Mezey, J. Gene expression network reconstruction by convex feature selection when incorporating genetic perturbations. PLoS Comput Biol 6, e1001014, doi:10.1371/journal.pcbi.1001014 (2010).

43 Heiman, M. G. & Walter, P. Prm1p, a pheromone-regulated multispanning membrane protein, facilitates plasma membrane fusion during yeast mating. J Cell Biol 151, 719–730, doi:10.1083/jcb.151.3.719 (2000).

44 Kibota, T. T. & Lynch, M. Estimate of the genomic mutation rate deleterious to overall fitness in E. coli. Nature 381, 694–696, doi:10.1038/381694a0 (1996).

45 Merlini, L., Dudin, O. & Martin, S. G. Mate and fuse: how yeast cells do it. Open Biol 3, 130008, doi:10.1098/rsob.130008 (2013).

46 Boocock, J., Sadhu, M. J., Durvasula, A., Bloom, J. S. & Kruglyak, L. Ancient balancing selection maintains incompatible versions of the galactose pathway in yeast. Science 371, 415–419, doi:10.1126/science.aba0542 (2021).

47 Nogi, Y., Matsumoto, K., Toh-e, A. & Oshima, Y. Interaction of super-repressible and dominant constitutive mutations for the synthesis of galactose pathway enzymes in Saccharomyces cerevisiae. Mol Gen Genet 152, 137–144, doi:10.1007/BF00268810 (1977).

48 Das Adhikari, A. K., Qureshi, M. T., Kar, R. K. & Bhat, P. J. Perturbation of the interaction between Gal4p and Gal80p of the Saccharomyces cerevisiae GAL switch results in altered responses to galactose and glucose. Mol Microbiol 94, 202–217, doi:10.1111/mmi.12757 (2014).

49 Dodd, D. M. B. Reproductive Isolation as a Consequence of Adaptive Divergence in Drosophila Pseudoobscura. Evolution 43, 1308–1311, doi:10.1111/j.1558-5646.1989.tb02577.x (1989).

50 Hurd, L. E., Eisenberg, R. M. Divergent selection for geotactic response and evolution of reproductive isolation in sympatric and allopatric populations of houseflies. American Naturalist 109, 353–358 (1975).

51 Barker, J. S. & Cummins, L. J. The Effect of Selection for Sternopleural Bristle Number on Mating Behaviour in DROSOPHILA MELANOGASTER. Genetics 61, 713–719, doi:10.1093/genetics/61.3.713 (1969).

52 van Dijken, F. R. & Scharloo, W. Divergent selection on locomotor activity in Drosophila melanogaster. II. Test for reproductive isolation between selected lines. Behav Genet 9, 555–561, doi:10.1007/BF01067351 (1979).

53 Brideau, N. J. et al. Two Dobzhansky-Muller genes interact to cause hybrid lethality in Drosophila. Science 314, 1292–1295, doi:10.1126/science.1133953 (2006).

54 Chen, J. et al. A triallelic system of S5 is a major regulator of the reproductive barrier and compatibility of indica-japonica hybrids in rice. Proc Natl Acad Sci U S A 105, 11436–11441, doi:10.1073/pnas.0804761105 (2008).

55 Lee, H. Y. et al. Incompatibility of nuclear and mitochondrial genomes causes hybrid sterility between two yeast species. Cell 135, 1065–1073, doi:10.1016/j.cell.2008.10.047 (2008).

56 Khan, A. I., Dinh, D. M., Schneider, D., Lenski, R. E. & Cooper, T. F. Negative epistasis between beneficial mutations in an evolving bacterial population. Science 332, 1193–1196, doi:10.1126/science.1203801 (2011).

57 Chen, P. & Zhang, J. Antagonistic pleiotropy conceals molecular adaptations in changing environments. Nat Ecol Evol 4, 461–469, doi:10.1038/s41559-020-1107-8 (2020).

58 Chen, P. & Zhang, J. Publisher Correction: Antagonistic pleiotropy conceals molecular adaptations in changing environments. Nat Ecol Evol 4, 488, doi:10.1038/s41559-020-1149-y (2020).

59 Sane, M., Miranda, J. J. & Agashe, D. Antagonistic pleiotropy for carbon use is rare in new mutations. Evolution 72, 2202–2213, doi:10.1111/evo.13569 (2018).

60 Blank, T. E., Woods, M. P., Lebo, C. M., Xin, P. & Hopper, J. E. Novel Gal3 proteins showing altered Gal80p binding cause constitutive transcription of Gal4p-activated genes in Saccharomyces cerevisiae. Mol Cell Biol 17, 2566–2575, doi:10.1128/MCB.17.5.2566 (1997).

61 Nishant, K. T. et al. The baker’s yeast diploid genome is remarkably stable in vegetative growth and meiosis. PLoS Genet 6, e1001109, doi:10.1371/journal.pgen.1001109 (2010).

62 Mumberg, D., Muller, R. & Funk, M. Yeast vectors for the controlled expression of heterologous proteins in different genetic backgrounds. Gene 156, 119–122, doi:10.1016/0378-1119(95)00037-7 (1995).

63 Salmeron, J. M., Jr., Leuther, K. K. & Johnston, S. A. GAL4 mutations that separate the transcriptional activation and GAL80-interactive functions of the yeast GAL4 protein. Genetics 125, 21–27, doi:10.1093/genetics/125.1.21 (1990).

64 Huxley, C., Green, E. D. & Dunham, I. Rapid assessment of S. cerevisiae mating type by PCR. Trends Genet 6, 236, doi:10.1016/0168-9525(90)90190-h (1990).

65 Bolger, A. M., Lohse, M. & Usadel, B. Trimmomatic: a flexible trimmer for Illumina sequence data. Bioinformatics 30, 2114–2120, doi:10.1093/bioinformatics/btu170 (2014).

66 Olson, N. D. et al. Best practices for evaluating single nucleotide variant calling methods for microbial genomics. Front Genet 6, 235, doi:10.3389/fgene.2015.00235 (2015).

