## Supplement Figures and Tables for "Rapid evolution of pre-zygotic reproductive barriers in allopatric populations"

**S1. Experimental design for adaptive evolution.**

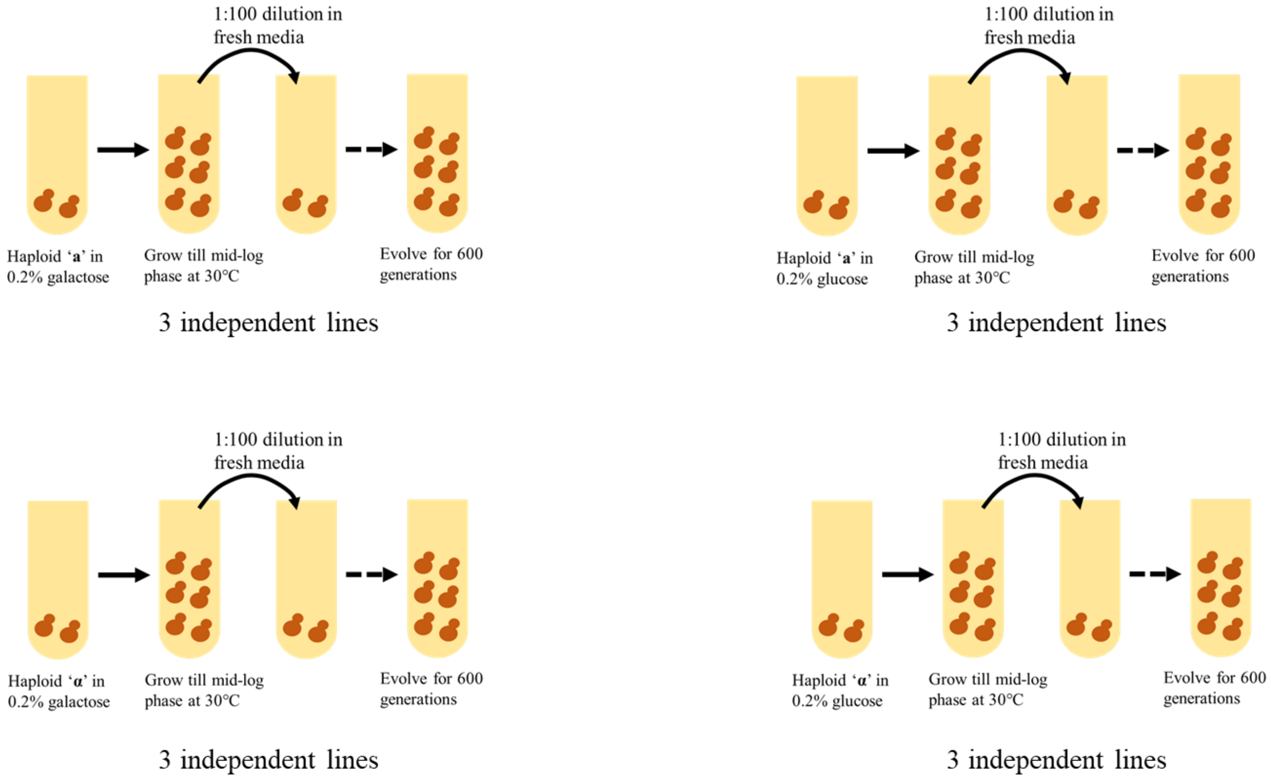


**Figure S1.** Twelve independent lines of haploid yeast were evolved in 0.2% glucose or 0.2% galactose, as shown above. Freezer stocks of the evolved lines were made every 200 generations.

**S2. Change in mating efficiency with adaptation in allopatry.**

**(A)** **(B)** **(C)**

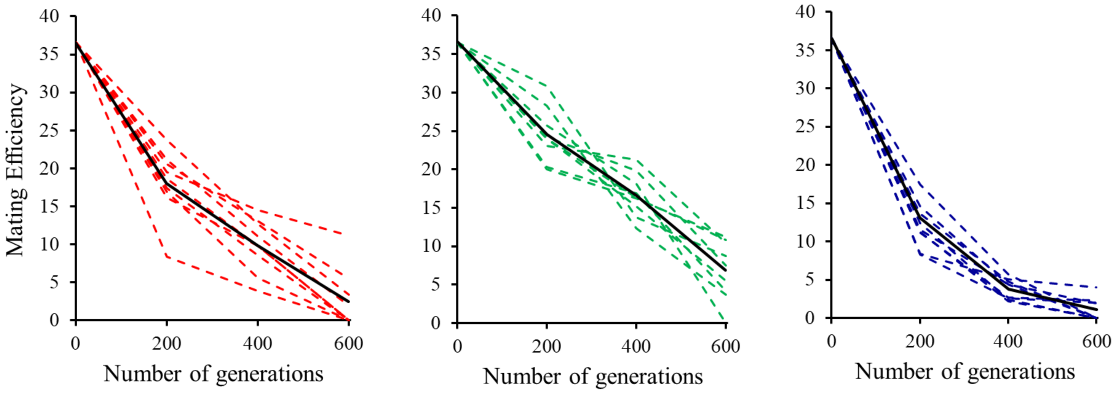


**(D)**

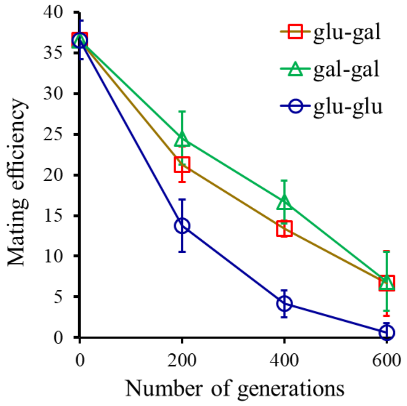


**Figure S2.** Mating efficiency between the **(A)** glucose- and galactose-evolved lines, **(B)** galactose- and galactose-evolved lines, and **(C)** glucose- and glucose-evolved lines at 200, 400, and 600 generations. Dotted lines indicate individually paired lines. Solid black line exhibits the average. **(D)** The mating efficiency data when lines glu1a and glu2a are excluded from the analysis. As shown in Figure S4, these lines underwent an autodiploidization event. All mating experiments were done in triplicate. Average is reported. Standard deviation of each is less than 10% of the data value for each mating experiment.

**S3. Two of the twelve evolved lines underwent an autodiploidization process.**

**(A)**

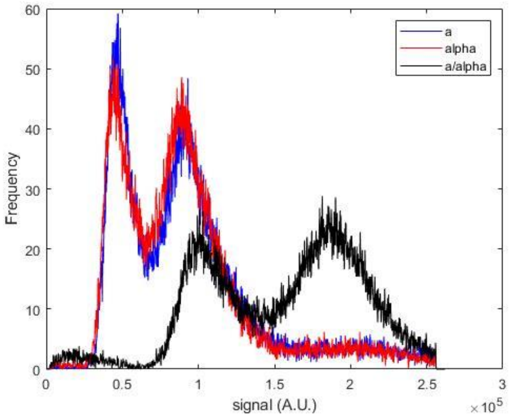


**(B)**

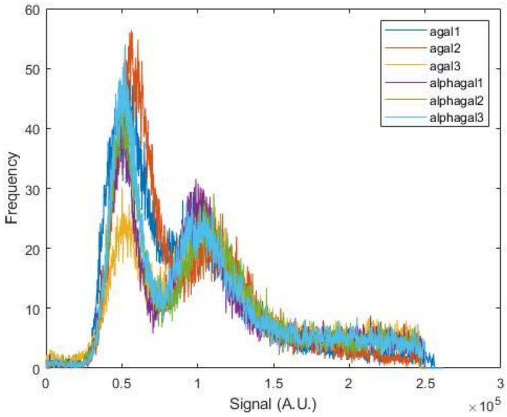


**(C)**

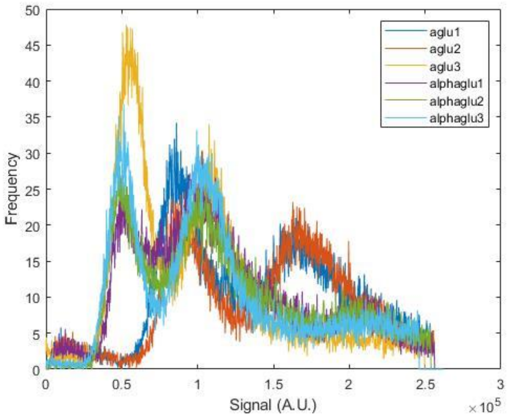


**Figure S3.** Two (glu1a and glu2a) out of the twelve evolved haploid lines underwent an autodiploidization event. **(A)** Ancestor a, α, and a/α, **(B)** the six lines evolved in galactose, and **(C)** the six lines evolved in glucose.

**S4. Hybrids generated from the evolved haploid lines.**

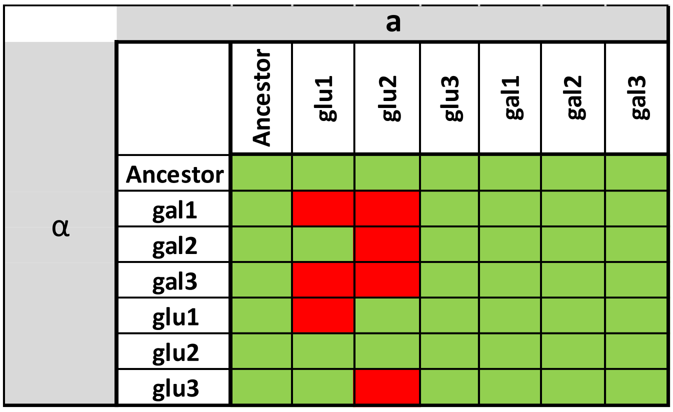


**Figure S4.** 42 out of the possible 49 hybrids were created (indicated in green). The cells highlighted in red refer to the hybrids, which could not be created.

**S5. Mitotic performance of the hybrids.**

**(A)**

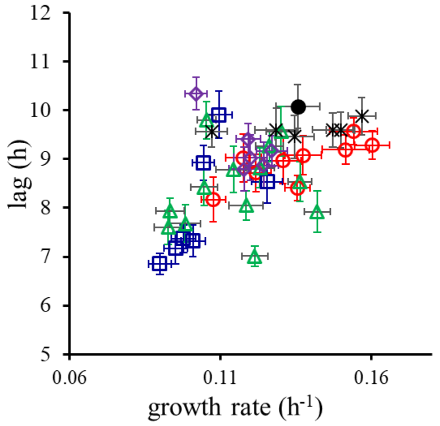


**(B)**

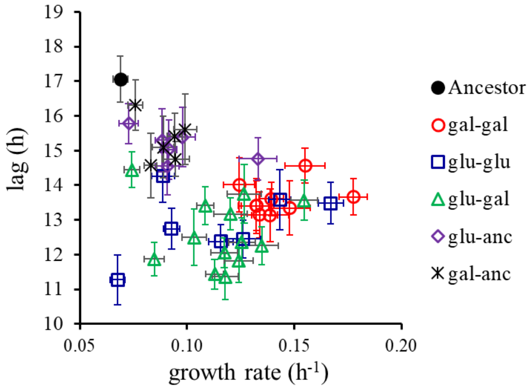


**(E)**

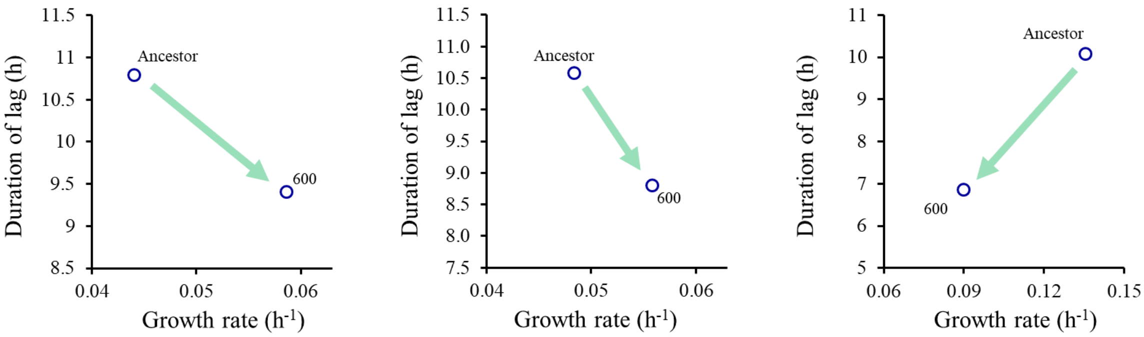


**Figure S5. Mitotic performance of the hybrids in glucose (A) and galactose (B).** Figure **(C)** shows Antagonism between beneficial mutations in glucose-adapted lines. Growth rate and lag phase duration in (Left) line glu1α, (Center) line glu3a, and (Right) glu1α-glu3a hybrid. The Ancestor comparisons are with ancestral **α** (left), ancestral **a** (center), and the ancestral **a**/**α** diploid (right). The number “600” represents the haploid at 600 generations (left and center) and the hybrid generated from these haploids (right). All experiments were done three independent times, and the average is represented. The standard deviation for each data point is less than 10% of the data value.

**S6. Comparison of the meiotic efficiency of the hybrids, compared to the ancestral diploid.**

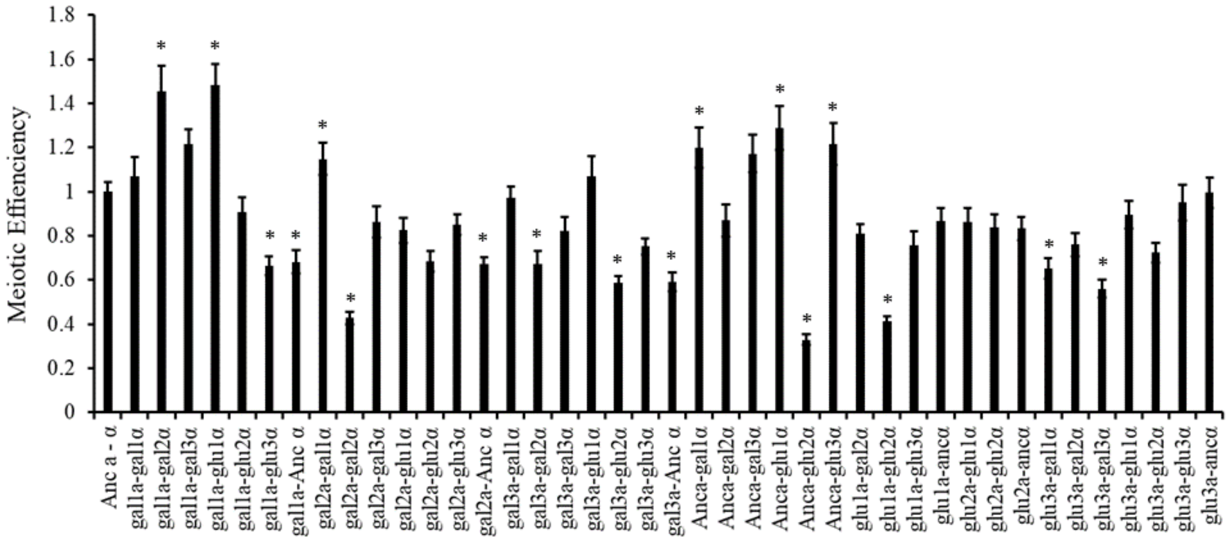


**Figure S6.** Meiotic efficiency of the hybrids, compared with the ancestor. All experiments were performed in triplicate. The average and standard deviation are represented. All hybrids which exhibit a meiotic efficiency which is statistically significantly different (p-value < 0.05) from that of the ancestor are indicated by *.

**S8. Spectrum of mutations in the galactose- and glucose-evolved haploid lines.**

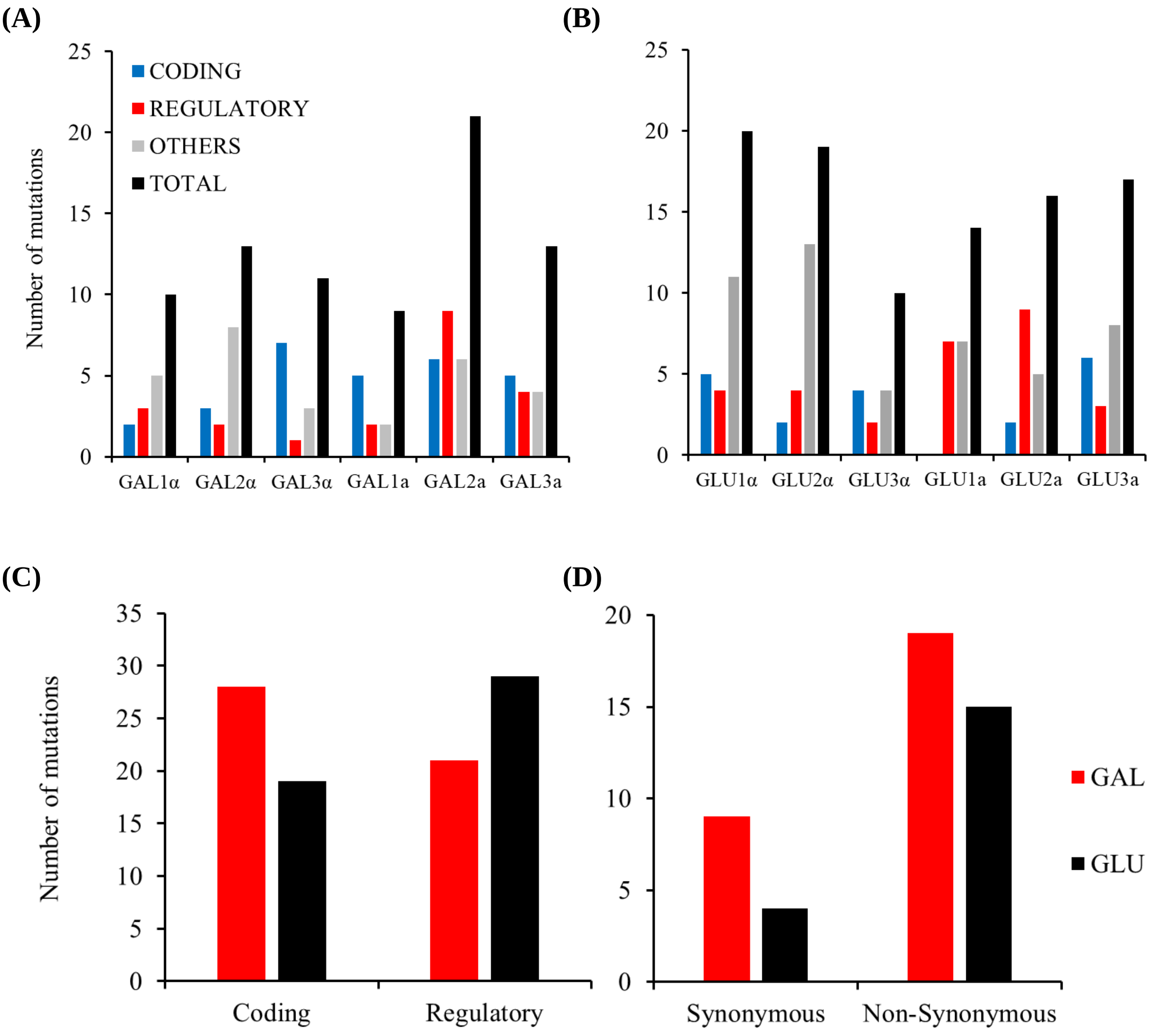


**Figure S8. (A)** and **(B)** Break-up of the nature of mutations in the 12 evolved lines. “Others” include SNPs and indels in intergenic regions of the chromosome. **(C)** and **(D)** Nature of mutations in the galactose- (red) and glucose-evolved (black) lines.

**S8. Relative fitness of the 44 haploid lines evolved under drift for 70 transfers in a mutation accumulation experiment.**

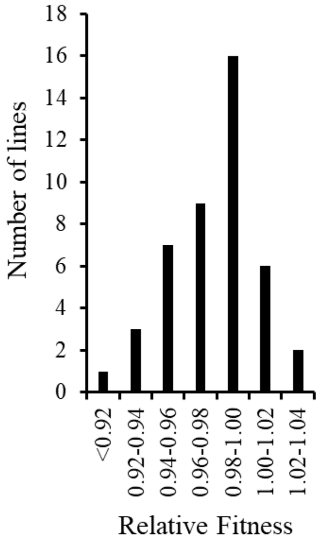


**Figure S8.** Distribution of mitotic fitness among the 44 mutation accumulation lines. The relative fitness on the x-axis represents the fitness of each line, with respect to its ancestral haploid fitness.

**S9. Hybrid mating efficiency, mitotic growth rate, and meiotic efficiency, as compared to the ancestor.**

**(A)** **(B)** **(C)**

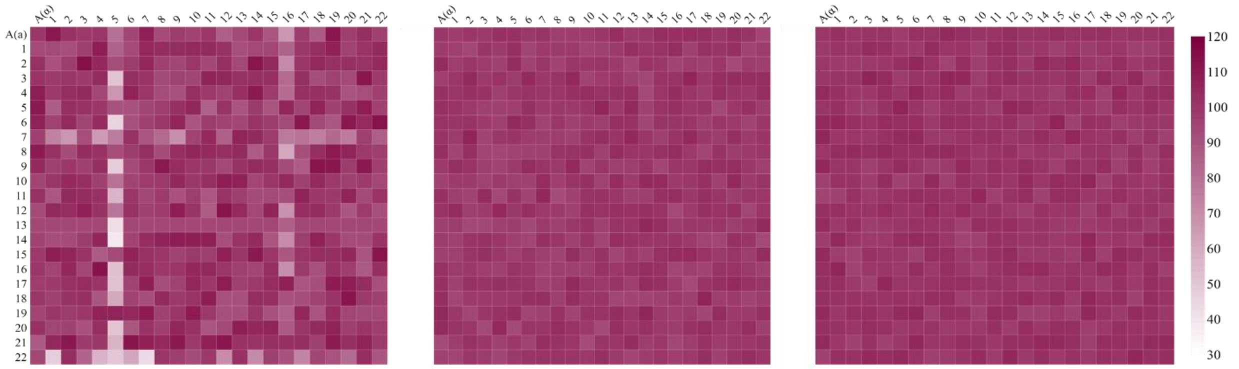


**Figure S9.** Pre-zygotic barriers arise faster than post-zygotic barriers. **(A)** Mating efficiency, **(B)** mitotic growth rate, and **(C)** meiotic efficiency of the hybrids formed from haploids evolved under drift in a mutation accumulation experiment.

**S10. Differences in mating barriers are strongly dictated by environment.**

**(A)** **(B)**

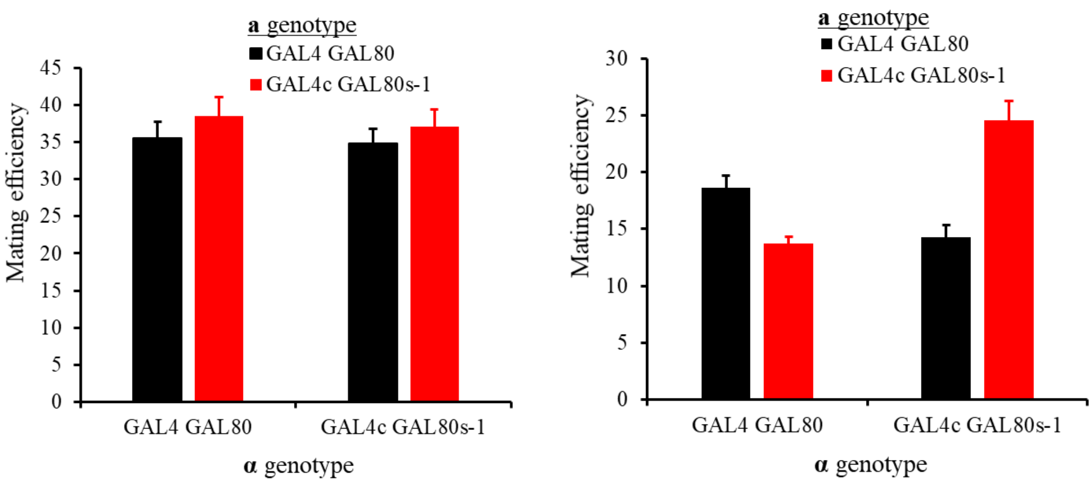


**Figure S10. SNPs can dictate mating efficiency. (A)** When grown in glucose, ancestor and the GAL4c GAL80s-1 mate with statistically identical efficiency. **(B)** When grown in melibiose, the mating efficiency between the ancestor haploids, the GAL4c GAL80s-1 haploids, and the ancestor-GAL4c GAL80s-1 haploids is statistically significantly different from each other.

**S11. Design for construction of the ancestral strains.**

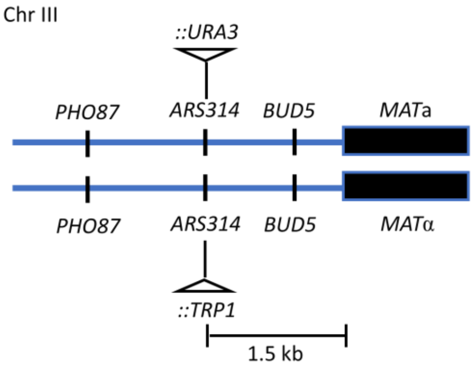


**Figure S11.** Markers URA3 and TRP1 inserted at the same site in ARS314 in haploids Mat α and a respectively, located between genes PHO87 and BUD5, approximately 1.5 kb from MAT locus. The insertion of markers does not disrupt either gene.

**S12. Experimental protocol for determination of mating efficiency.**

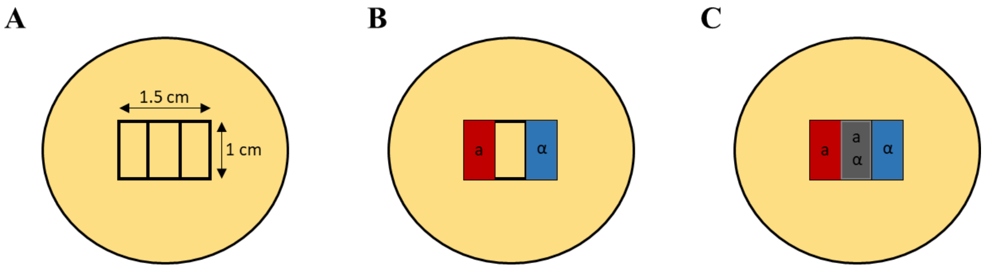


**Figure S12. Mating Efficiency Protocol.** A fresh YPD plate was marked as shown in **(A)**. A monolayer of the two haploids (**a** and **α**) was laid on the two rectangular areas, as shown in **B**. The monolayer was then allowed to grow at 30 °C for 24 hours. An equal number of **a** and **α** haploids from the red and blue marked areas in the figure were suspended in PBS buffer and mixed well. A monolayer of this mix of the two haploids was laid in the center rectangle of the marked area. The **a** and **α** mix was allowed to mate for 7 hours at 30 °C. Cells from the center rectangle were then picked up and spread on a YPD plate for single colonies. For all mating efficiencies, at least 500 colonies were analyzed for each experiment. All experiments were done three independent times. For all mating experiments, except Fig. S10B, cells grown in YPD were used to lay in a monolayer for (B). For the mating experiment in Fig. S10B, cells were grown to mid-log phase in 2% melibiose, and then laid down on a YPD plate, as shown in Fig. S12B.

**Table S1.**

**Table S1. p-value for the growth rate and lag phase duration for the hybrids in glucose.** (all values less than 0.05 are indicated in red)

| **Growth Rate** | Ancestor | gal-gal hybrid | glu-glu hybrid | glu-gal hybrid | glu-ancestor hybrid |
|---|---|---|---|---|---|
| gal-gal hybrid | 0.9400 | | | | |
| glu-glu hybrid | 0.0026 | 0.0016 | | | |
| glu-gal hybrid | 0.0169 | 0.0119 | 0.0252 | | |
| glu-ancestor hybrid | 0.0264 | 0.0192 | 0.0144 | 0.5567 | |
| gal-ancestor hybrid | 0.7669 | 0.6807 | 0.0011 | 0.0078 | 0.0121 |

| **Lag phase duration** | Ancestor | gal-gal hybrid | glu-glu hybrid | glu-gal hybrid | glu-ancestor hybrid |
|---|---|---|---|---|---|
| gal-gal hybrid | 0.0273 | | | | |
| glu-glu hybrid | 0.0032 | 0.0366 | | | |
| glu-gal hybrid | 0.0078 | 0.1658 | 0.2495 | | |
| glu-ancestor hybrid | 0.0846 | 0.3674 | 0.0171 | 0.0599 | |
| gal-ancestor hybrid | 0.2482 | 0.0944 | 0.0061 | 0.0183 | 0.3524 |

**Table S2.**

**Table S2. p-value for the growth rate and lag phase duration for the hybrids in galactose.**

| **Growth Rate** | Ancestor | gal-gal hybrid | glu-glu hybrid | glu-gal hybrid | glu-ancestor hybrid |
|---|---|---|---|---|---|
| gal-gal hybrid | 0.0001 | | | | |
| glu-glu hybrid | 0.0003 | 0.0077 | | | |
| glu-gal hybrid | 0.0003 | 0.0102 | 0.7981 | | |
| glu-ancestor hybrid | 0.0021 | 0.0012 | 0.0145 | 0.0136 | |
| gal-ancestor hybrid | 0.0035 | 0.0006 | 0.0038 | 0.0046 | 0.1985 |

| **Lag phase duration** | Ancestor | gal-gal hybrid | glu-glu hybrid | glu-gal hybrid | glu-ancestor hybrid |
|---|---|---|---|---|---|
| gal-gal hybrid | 0.0031 | | | | |
| glu-glu hybrid | 0.0015 | 0.2694 | | | |
| glu-gal hybrid | 0.0010 | 0.1377 | 0.6340 | | |
| glu-ancestor hybrid | 0.0311 | 0.0609 | 0.0192 | 0.0116 | |
| gal-ancestor hybrid | 0.0425 | 0.0487 | 0.0163 | 0.0100 | 0.8192 |

**Table S3.**

**Table S3. List of mutations in the evolved lines.** Genes in black text have mutations in the coding region. Genes in bold blue have mutations in the promoter regions. Intergenic mutations are not indicated in the tables below. See Annexure I for more details.

| Gala1 | Gala2 | Gala3 | Galα1 | Galα2 | Galα3 |
|---|---|---|---|---|---|
| MNL1 | SRD1 | PMD1 | COQ4 | MNS1 | CRD1 |
| FLO1 | CCH1 | YGL260W | OPY2 | MRT4 | PRM7 |
| RTT105 | MSS11 | ZAP1 | **CWC23** | Hyp. | LAM4 |
| YIL163C | POP2 | YPL277C | **PSR1** | **RIB7** | MNL1 |
| CCC1 | TMA23 | BRO1 | **BFA1** | **PRO2** | GAL2 |
| **LHP1** | SAP190 | **YNCB0008W** | | **RRM3** | KIN2 |
| **DET1** | **PNC1** | **GYP6** | | **ERC1** | YLR406C-A |
| | **AMD1** | **CYC1** | | | YNR065C |
| | **FRT1** | **CDC45** | | | |
| | **NOC4** | | | | |
| | **BOR1** | | | | |
| | **SDA1** | | | | |
| | **tE(UUC)I** | | | | |
| | **SPT8** | | | | |
| | **VTC5** | | | | |
| **Glua1** | **Glua2** | **Glua3** | **Gluα1** | **Gluα2** | **Gluα3** |
| **COS111** | FIT1 | PRM7 | ABP1 | SDH1 | LAA1 |
| **PCL6** | BIN1 | RTG2 | RPO21 | HSP104 | MNS1 |
| **ERC1** | **DAD3** | FAR1 | YHL008C | **KAR4** | BUL2 |
| **YPT52** | **YIH1** | ARG3 | MNN4 | **RPL8A** | UBP7 |
| **YKR075C** | **XBP1** | AVL9 | PAN3 | **SUF8** | **KAP95** |
| **YLR053C** | **TGL5** | MNS1 | **PRP11** | **SLY41** | **MRS6** |
| **RPA43** | **YOR381W-A** | **ASM4** | **MGA1** | | |
| | **AMD1** | **VHS1** | **YJL132W** | | |
| | **TAL1** | **SLT2** | **FRE7** | | |
| | **CBF1** | | | | |
| | **YFH7** | | | | |

**Table S4.**

**Table S4. List of primers used in this study**

| Name | Sequence (5' – 3') |
|---|---|
| pSc011 | GTT GAC TGT AAT ATC TGT AAA AGA TTA CAT CTA ATT TAC GTT CAA TTC AAT TCA TCA TTT |
| pSc012 | AGA TGA CTT CCT TTT GCT TCT TGT ACG CTC ACA AAT AAT TTT AGT TTT GCT GGC CGC ATC |
| pSc014 | GTT GAC TGT AAT ATC TGT AAA AGA TTA CAT CTA ATT TAC GAA CGA CAT TAC TAT ATA TAT |
| pSc015 | AGA TGA CTT CCT TTT GCT TCT TGT ACG CTC ACA AAT AAT TCT ATT TCT TAG CAT TTT TGA |
| pSc018 | GAA AGA GGC GGT AGC CTA AAG ATA CGG TAA TTG AAA CGT TTC CTA TGC ACA ATC TTA AAC CTT TTT AGG TAA TTG ATT AAG TTG ACT GTA ATA TCT GTA AAA GAT TAC ATC TAA TTT ACG |
| pSc019 | AAC GAC AAC AAA GAC AGG GCG TAT CAA GTC AGT ATA GCT AAG GTT CCA AGG CTT ACC TAA AAA CAG AAC TGT TCA AAG AAA GAT GAC TTC CTT TTG CTT CTT GTA CGC TCA CAA ATA ATT |
| pSc020 | ATT TTT CAA TGC ATC GGA TTA CTT TTC CCA CGT GCG AAA TCA TCA ATT AAT TAG ATT GAA AAA AGG GTA AGG GAA AAT AAG AAA GAG GCG GTA GCC TAA AGA TAC GGT AAT TGA AAC GTT |
| pSc021 | AGC AAT ATG AAG ATA TAC ACT GTT CAA ATA CTA CTG CAA GAA GCT TAT TAG GGG CTA TGA TAA AGG TGC ACA CTT TAT ATA ACG ACA ACA AAG ACA GGG CGT ATC AAG TCA GTA TAG CTA |

**Annexure I. List of mutations in the 12 evolved lines.**

| Gal1a | | |
|---|---|---|
| | **Mutation.** | **Gene function. Gene function and adaptation. Gene function and mating behavior.** |
| 1 | I/207330 [(missense_variant c.3928G>A p.Val1310Ile) in FLO1] | FLOcculation, Lectin-like protein involved in flocculation, cell wall protein that binds mannose chains on the surface of other cells [1-3]. Null mutants show shows reduced competitive fitness in minimal medium [4]. Overexpression causes increase in mating efficiency [5]. |
| 2 | V/366862 [(missense_variant c.61G>A p.Ala21Thr) in RTT105] | Regulator of Ty1 Transposition, Chaperone for Replication Protein A complex (RPA), involved in nuclear import of RPA and its binding to ssDNA at replication forks; has a role in regulation of Ty1 transposition [6,7]. |
| 3 | VIII/508350 [(stop_gained c.2032G>T p.Glu678*) in MNL1] | MaNnosidase-Like protein, Alpha-1,2-specific exomannosidase of the endoplasmic reticulum [8]. Sporulation efficiency increased in null mutants [9]. |
| 4 | IX/37201 [(frameshift_variant c.48_51dupAAAA p.Val18fs) in YIL163C] | |
| 5 | XII/577545 [(missense_variant c.721G>T p.Gly241Cys) in CCC1] | Cross-Complements Ca(2+) phenotype of csg1, Vacuolar Fe2+/Mn2+ transporter, suppresses respiratory deficit of yfh1 mutants [10-13]. |
| 6 | IV/363894 [C to T 58 bp upstream of LHP1] | La-Homologous Protein, RNA binding protein required for maturation of tRNA and U6 snRNA, acts as a molecular chaperone for RNAs transcribed by polymerase III [14-16]. Overexpression causes decrease in vegetative growth [17,18]. |
| 7 | IV/558164 [C to CT 104 bp upstream of DET1] | Decreased Ergosterol Transport, Acid phosphatase; involved in the non-vesicular transport of sterols in both directions between the endoplasmic reticulum and plasma membrane [19-21]. |
| 8 | X/745722 [(intergenic_region n.745722_745723insGGT)] | |
| 9 | XI/469215 [C to A] | |
| 10 | XII/306619 [ATA to T] | |
| **Gal2a** | | |
| | **Mutation.** | **Gene function. Gene function and adaptation. Gene function and mating behavior.** |
| 1 | III/148500 [(missense_variant c.404G>A p.Arg135Gln) in SRD1] | Protein involved in the processing of pre-rRNA to mature rRNA, contains a C2/C2 zinc finger motif, srd1 mutation suppresses defects caused by the rrp1-1 mutation [22]. |
| 2 | VII/926639 [(synonymous_variant c.1944C>A p.Pro648Pro) in CCH1] | Calcium Channel Homolog, Voltage-gated high-affinity calcium channel; involved in calcium influx in response to some environmental stresses as well as exposure to mating pheromones [23,24]. Calcium levels are linked with glycolysis [25,26]. CCH1 gene is involved in calcium influx and mating [27,28]. |
| 3 | XIII/588585 [(synonymous_variant c.966A>G p.Gln322Gln) in MSS11] | Multicopy Suppressor of STA genes, Transcription factor, involved in regulation of invasive growth and starch degradation, controls the activation of FLO11 and STA2 in response to nutritional signals, forms a heterodimer with Flo8p that interacts with the Swi/Snf complex during transcriptional activation of FLO1, FLO11, and STA1 [29-32]. Mss11p is a transcription factor regulating pseudohyphal differentiation, invasive growth and starch metabolism in Saccharomyces cerevisiae in response to nutrient availability [31]. |
| 4 | XIV/720276 [(disruptive_inframe_deletion c.357_371del ACAACAGCAGCAACA p.Gln120_Gln124del) in POP2] | PGK promoter directed OverProduction, Subunit of Ccr4-Not complex that mediates 3' to 5' mRNA deadenylation [33,34]. Required for glucose-derepression of gene expression [35]. Decreased sporulation efficiency in null mutants [9,36,37]. |
| 5 | XIII/804970 [(missense_variant c.515G>A p.Gly172Glu) in TMA23] | Translation Machinery Associated, Nucleolar protein implicated in ribosome biogenesis, deletion extends chronological lifespan [38-40]. |
| 6 | XI/495887 [(conservative_inframe_deletion c.1632_1633delCAA p.Val544_Glu545insGln) in SAP190] | Sit4 Associated Protein, Protein that forms a complex with the Sit4p protein phosphatase, required for Sit4p function [41]. |
| 7 | IV/621879 [GAAA to G 233 bp upstream of VTC5] | Vacuole Transporter Chaperone, Novel subunit of the vacuolar transporter chaperone complex, vacuolar transmembrane protein that regulates biosynthesis of polyphosphate [42-44]. |
| 8 | VII/982274 [CTG to AAAAAACTA 206 bp upstream of SDA1] | Severe Depolymerization of Actin, Protein required for actin organization and passage through Start; highly conserved nuclear protein, required for actin cytoskeleton organization, plays a critical role in G1 events, involved in 60S ribosome biogenesis [45-47]. Growth in exponential phase is decreased due to overexpression [45]. |
| 9 | IX/370424 [T to C (intergenic_region n.370424T>C ) tE(UUC)I] | |

| | | |
|---|---|---|
| 10 | XII/253093 [G to GAA 13 bp upstream of SPT8] | SuPpressor of Ty, Subunit of the SAGA transcriptional regulatory complex, not present in SAGA-like complex SLIK/SALSA, required for SAGA-mediated inhibition at some promoters [48]. Decreased sporulation efficiency in null mutants [49]. |
| 11 | XV/925072 [C to CA 32 bp upstream of FRT1] | Functionally Related to TCP1, Tail-anchored ER membrane protein, promotes cell growth in stress conditions, possibly via a role in posttranslational translocation [50-53]. |
| 12 | XVI/821526 [CT to TTC 103 bp upstream of NOC4] | NucleOlar Complex associated , Nucleolar protein; forms a complex with Nop14p that mediates maturation and nuclear export of 40S ribosomal subunits, relocalizes to the cytosol in response to hypoxia [54-56]. |
| 13 | XIV/119025 [T to C 243bp upstream of BOR1] | BORon transporter, Boron efflux transporter of the plasma membrane [57]. Null mutant displays a decreased ability to utilize galactose as carbon source and both arginine and glutamate as nitrogen sources [58]. |
| 14 | XIII/208877 [T to C 17 bp upstream of AMD1] | AMP Deaminase, tetrameric enzyme that catalyzes the deamination of AMP to form IMP and ammonia, thought to be involved in regulation of intracellular purine (adenine, guanine, and inosine) nucleotide pools [59]. |
| 15 | VIII/382367 [TA to T Intron RPL42B] | Ribosomal Protein of the Large subunit, Ribosomal 60S subunit protein L42B [60]. |
| 16 | VII/428071 [AAT to A 124 bp upstream of PNC1] | Pyrazinamidase and NiCotinamidase, Nicotinamidase that converts nicotinamide to nicotinic acid; part of the NAD(+) salvage pathway, required for life span extension by calorie restriction [61,62]. |
| 17 | X/470124 [ATAATAG to A] | |
| 18 | X/470135 [A to G] | |
| 19 | X/470148 [A to G] | |
| 20 | XII/14561 [A to AT] | |
| 21 | X/177910 [G to GAT] | |
| | **Gal3a** | |
| | **Mutation.** | **Gene function. Gene function and adaptation. Gene function and mating behavior.** |
| 1 | V/427351 [(missense_variant c.3099C>A p.Asp1033Glu) in PMD1] | Paralog of MDS3, Protein with an N-terminal kelch-like domain, putative negative regulator of early meiotic gene expression [63]. Putative negative regulator of early meiotic gene expression [64]. |
| 2 | VII/7008 [(missense_variant c.149C>A p.Ser50Tyr) in YGL260W] | |
| 3 | X/332667 [(disruptive inframe deletion c.359_406del CGTCA TTAAC AAAAT ATAAT GATAC TGCAA CGTAT AATTC TAATA ATC p.Pro120_Asn135del) in ZAP1] | Zinc-responsive Activator Protein, Zinc-regulated transcription factor, binds to zinc-responsive promoters to induce transcription of certain genes in presence of zinc [65,66]. |
| 4 | XVI/16500 [(synonymous_variant c.369T>C p.His123His) in YPL277C] | |
| 5 | XVI/395414 [(missense_variant c.1377G>A p.Met459Ile) in BRO1] | BCK1-like Resistance to Osmotic shock, Cytoplasmic class E vacuolar protein sorting (VPS) factor, coordinates protein sorting and deubiquitination in the multivesicular body (MVB) pathway by recruiting Doa4p to endosomes [67-70]. Null mutant has a sporulation defect [9]. |
| 6 | [C to G 13bp downstream of YNCB0008W] | lncRNA antisense to GAL10 and overlapping GAL1 mRNAs; GAL10-ncRNA transcription recruits Set2p methyltransferase and histone deacetylation activities in cis, leading to stable changes in chromatin structure; acts to enhance glucose repression of GAL1-10 induction at low environmental sugar concentrations; expression driven by Reb1p; does not appear to control GAL1 induction [71,72]. |
| 7 | X/359824 [GA to G 148 bp upstream of GYP6] | Gtpase-activating protein of Ypt6 Protein, GTPase-activating protein (GAP) for yeast Rab family member Ypt6p, involved in vesicle mediated protein transport [73,74]. |
| 8 | X/525782 [TAAAAA to T, 401 bp upstream of ANB1, & 882 bp upstream of CYC1] | CYtochrome C, electron carrier of mitochondrial intermembrane space that transfers electrons from ubiquinone-cytochrome c oxidoreductase to cytochrome c oxidase during cellular respiration [75,76]. |
| 9 | XVI/17926 [G to A 22 Bp upstream of YPL276W] | |
| 10 | XII/346105 [GA to G 164 bp upstream of CDC45] | Cell Division Cycle , DNA replication initiation factor [77,78]. |
| 11 | IV/164943 [C to CT] | |
| 12 | XVI/64363 [A to AACACCAGTTTCTTTGAGG] | |
| | **Gal1α** | |
| | **Mutation.** | **Gene function. Gene function and adaptation. Gene function and mating behavior.** |
| 1 | IV/858770 [(Missense variant c.634T>G p.Phe212Val) in COQ4] | Protein with a role in ubiquinone (Coenzyme Q) biosynthesis [79-81]. Respiratory growth is absent in null mutants [82]. |
| 2 | XVI/696343 [(disruptive_inframe_insertion c.462_476dup | Overproduction-induced Pheromone-resistant Yeast, Integral membrane protein that acts as a membrane anchor for Ste50p, and as a regulator of the filamentous growth pathway. Overproduction blocks cell cycle arrest in the presence of mating pheromone, relocalizes from vacuole to plasma membrane |

| | | |
|---|---|---|
| | AGACGATGAGGATGA p.Glu154_Asp158dup) in OPY2] | upon DNA replication stress [83,84]. MAPK Pathway responds to glucose starvation through Mig1/2 [85]. Null mutants display distal-unipolar budding defect under filamentous growth-inducing conditions [85]. |
| 3 | VII/270311 [A to ATG 167 bp upstream of CWC23] | Component of a complex containing Cef1p; putatively involved in pre-mRNA splicing [86-88]. Abnormal sporulation efficiency in null mutants [89]. |
| 4 | XII/130916 [CC to TTTCG 303 bp upstream of PSR1] | Plasma membrane Sodium Response, Plasma membrane-associated protein phosphatase [90]. Required along with binding partner Msn2p for inhibition of TORC1 in response to limiting amino acids [91,92]. |
| 5 | X/533814 & 533833 [G to GT 213, and C to T 194 bases upstream of BFA1] | Byr-Four-Alike, Subunit of a two-component GTPase-activating protein, Bfa1p-Bub2p, contributes to GAP activity, inactivating Tem1 by stimulating GTP hydrolysis following damage or misalignment of the mitotic spindle also functions as a guanine-nucleotide exchange inhibitor (GDI) for Tem1p.Involved in multiple cell cycle checkpoint pathways that control mitotic exit [93-97]. Regulated by FKH1 during maintenance of stationary phase in response to starvation [98]. The GAP complex Bfa1-Bub2 are dispensable for activation of the kinases Cdc15 and Dbf2/20-Mob1, as well as spore formation [99,100]. |
| 6 | IV/344004 [G to GTA] | |
| 7 | XII/172018 [AATATATAT to A] | |
| 8 | XIII/224327 [AT to A] | |
| 9 | XIII/224338 [CG to AA] | |
| 10 | XIII/234752 [A to AT] | |
| | **Gal2α** | |
| | **Mutation.** | **Gene function. Gene function and adaptation. Gene function and mating behavior.** |
| 1 | X/669207 [(missense_variant c.1564G>T p.Val522Phe) in MNS1] | Alpha-1,2-mannosidase; involved in ER-associated protein degradation (ERAD), catalyzes the removal of one mannose residue from a glycosylated protein, converting the modification from Man9GlcNAc to Man8GlcNAc. Catalyzes the last step in glycoprotein maturation in the ER and is critical for ER protein degradation [101-104]. |
| 2 | XI/426930 [(missense_variant c.689G>A p.Ser230Asn) in MRT4] | mRNA Turnover 4, protein involved in mRNA turnover and ribosome assembly, required at post-transcriptional step for efficient retrotransposition, localizes to the nucleolus [105,106]. |
| 3 | XIV/751343 [(synonymous_variant c.2358C>T p.Ile786Ile) in Hypothetical Protein] | |
| 4 | II/547397 [G to A 63 bp upstream of RIB7] | RIBoflavin biosynthesis, catalyzes the second step of the riboflavin biosynthesis pathway [107-109]. |
| 5 | VIII/173117 [ATGAAAAAAAA AAAAATAATA to TGAAAAAAA AAATAAT 172 bp upstream of RRM3 and 227 bp upstream of ERC1] | rDNA Recombination Mutation, DNA helicase involved in rDNA replication [110]. Ethionine resistance conferring [111,112]. Overexpression causes abnormal budding [17]. |
| 6 | XV/923126 [G to GAT 221 bp upstream of PRO2] | PROline requiring, Gamma-glutamyl phosphate reductase, catalyzes the second step in proline biosynthesis [113]. |
| 7 | IV/691402 [T to ATATATA] | |
| 8 | VI/157816 [AT to A] | |
| 9 | VI/157828 [A to ATCT] | |
| 10 | VIII/370543 [AAT to A] | |
| 11 | IX/51679 [CATTATT to C] | |
| 12 | X/358212 [GTTT to T] | |
| 13 | X/654573 [GAA to G] | |
| 14 | XII/373555 [TTG to T] | |
| | **Gal3α** | |
| | **Mutation.** | **Gene function. Gene function and adaptation. Gene function and mating behavior.** |
| 1 | IV/202356 [(missense_variant c.215A>G p.Asn72Ser) in CRD1] | CaRDiolipin synthase,produces cardiolipin, which is a phospholipid of the mitochondrial inner membrane that is required for normal mitochondrial membrane potential and function and for correct integration of membrane-multispanning proteins into the mitochondrial outer membrane, required to |

| | | |
|---|---|---|
| | | maintain tubular mitochondrial morphology and functions in mitochondrial fusion, also required for normal vacuolar ion homeostasis [114-117]. |
| 2 | IV/383938 [(synonymous_variant c.144C>T p.Thr48Thr) in PRM7] | Pheromone-Regulated Membrane protein. |
| 3 | VIII/263773 [(synonymous_variant c.3066A>G p.Gly1022Gly) in LAM4] | Lipid transfer protein Anchored at Membrane contact sites , Sterol-binding protein that localizes to puncta in the cortical ER [118]. Deletion of LAM genes inhibits the regulated death of Saccharomyces cerevisiae yeast cells induced by the mating pheromone [119]. |
| 4 | VIII/507586 [(missense_variant c.1268A>T p.Asp423Val) in MNL1] | MaNnosidase-Like protein, Alpha-1,2-specific exomannosidase of the endoplasmic reticulum [8]. Sporulation efficiency is increased in null mutants [9]. |
| 5 | XII/290859 [(synonymous_variant c.648G>C p.Leu216Leu) in GAL2] | Galactose permease, required for utilization of galactose, also able to transport glucose [120-123]. |
| 6 | XII/333094 [(synonymous_variant c.505A>C p.Arg169Arg) in KIN2] | S/T protein kinase, regulates polarized exocytosis and the Ire1p-mediated UPR, regulates HAC1 mRNA translocation, splicing and translation with KIN1 during ER stress [124-127]. |
| 7 | XII/932282 [(frameshift_variant c.64_73delAGTATACATC p.Ser22fs) in YLR406C-A] | |
| 8 | XIV/751807 [(missense_variant c.1894C>T p.Pro632Ser) in YNR065C] | |
| 9 | VI/270133 [TGGGTGT to GGTGTGTG] | |
| 10 | IX/105805 [TA to T] | |
| 11 | IX/105811 [TA to T] | |
| | **Glu1a** | |
| | **Mutation.** | **Gene function. Gene function and adaptation. Gene function and mating behavior.** |
| 1 | II/628892 [TATAAG to ATATAA GCATA276 bp upstream of COS111] | Ciclopirox Olamine Sensitive, Protein required for antifungal drug ciclopirox olamine resistance [128]. Overexpression causes abnormal cell morphology and budding [17]. |
| 2 | V/272558 [GT to T 66 bp upstream of PCL6] | Pho85 CycLin, Pho85p cyclin of the Pho80p subfamily, forms the major Glc8p kinase together with Pcl7p and Pho85p, involved in the control of glycogen storage by Pho85p [129]. |
| 3 | VIII/173184 [G to GA 160 bp upstream of ERC1] | Ethionine Resistance Conferring, Member of the multi-drug and toxin extrusion (MATE) family. |
| 4 | XI/465172 [G to GTATA 195 bp downstream of YPT52] | Yeast Protein Two, Endosomal Rab family GTPase, required for vacuolar protein sorting, endocytosis and multivesicular body (MVB) biogenesis and sorting [130]. Null mutants show decreased utilization of galactose as carbon source [58]. |
| 5 | XI/580897 [ATATG to A 147 bp upstream of YKR075C] | |
| 6 | XII/248700 [A to ATT 274 bp upstream of YLR053C] | |
| 7 | XV/960346 [A to C 164 bp upstream of RPA43] | RNA polymerase I subunit A43 [131], exhibits a decreased competitive fitness in a null mutant decreased [4]. |
| 8 | II/40582 [GATATATAC to G] | |
| 9 | II/40600 [T to C] | |
| 10 | XI/638422 [AT to A] | |
| 11 | XII/289370 [TGGG to GGA] | |
| 12 | XVI/711293 [C to CT] | |
| 13 | XII/373658 [CTCGTGGA CGTGGAC to TTCGTGGAT] | |
| | **Glu2a** | |
| | **Mutation.** | **Gene function. Gene function and adaptation. Gene function and mating behavior.** |
| 1 | IV/1504389 [(missense_variant c.512C>T p.Thr171Ile) in FIT1] | Mannoprotein incorporated into the cell wall, incorporated via a glycosylphosphatidylinositol (GPI) anchor [132,133]. |
| 2 | XIV/131661 [(conservative_inframe_ insertion& synonymous_variant c.3721_3723delCCGins TCGCCTCCTCCT p.Pro1240_Pro1241ins SerProPro) in BNI1] | Bud Neck Involved, Formin, polarisome component, involved in cell processes such as budding and mitotic spindle orientation which require the formation of polarized actin cables [134-136]. Null mutant exhibits decreased mating efficiency [137,138], abnormal mating projection [139], decreased shmoo formation [137], and abnormal budding pattern [140]. |

| | | |
|---|---|---|
| 3 | II/684840 [A to AT 137 bp upstream of DAD3] | Essential subunit of the Dam1 complex (aka DASH complex), complex couples kinetochores to the force produced by MT depolymerization thereby aiding in chromosome segregation, is transferred to the kinetochore prior to mitosis [141,142]. |
| 4 | III/224251 [C to CTT 21 bp upstream of YIH1] | Negative regulator of eIF2 kinase Gcn2p, regulation of translation in response to starvation via regulation of Gcn2p [143,144]. Yih1 interacts with the Cyclin Dependent Kinase Cdc28 and promotes cell cycle progression through G2/M in budding yeast [145]. |
| 5 | IX/177463 [GCA to G 213 bp upstream of XBP1] | XhoI site-Binding Protein, Transcriptional repressor; binds promoter sequences of cyclin genes, CYS3, and SMF2; not expressed during log phase of growth, but induced by stress or starvation during mitosis, and late in meiosis [146-148]. Decreased sporulation efficiency in null mutants[147], and budding index abnormal budding index in overexpression [17]. |
| 6 | XV/479293 [GTTTTAT to TGTTTTAC 105 bp upstream of TGL5] | Bifunctional triacylglycerol lipase and LPA acyltransferase,involved in triacylglycerol mobilization, catalyzes acylation of lysophosphatidic acid (LPA), potential Cdc28p substrate [149]. |
| 7 | XV/1058395 [ATTA to TT 28 bp upstream of YOR381W-A] | |
| 8 | XIII/208862 [AT to A 2 bp upstream of AMD1] | AMP deaminase;tetrameric enzyme that catalyzes the deamination of AMP to form IMP and ammonia [59,150,151]. Null mutant exhibits a decreased growth rate on rich media and decreased competitive fitness on both rich and synthetic complete media, null mutant displays a delay in acceleration of budding after a glucose pulse [152]. Null mutant displays a delay in acceleration of budding after a glucose pulse, budding delayed [153]. |
| 9 | XII/837543 [A to G 186 bp upstream of TAL1] | Transaldolase, enzyme in the non-oxidative pentose phosphate pathway, converts sedoheptulose 7-phosphate and glyceraldehyde 3-phosphate to erythrose 4-phosphate and fructose 6-phosphate [53,154]. Overexpression causes increased rate of utilization of carbon source [155]. |
| 10 | X/548539 [C to CA 220 bp upstream of CBF1] | Basic helix-loop-helix (bHLH) protein, associates with kinetochore proteins, required for chromosome segregation, protein abundance increases in response to DNA replication stress [156-159]. |
| 11 | VI/159212 [GA to G 87 bp upstream of YFH7] | Putative kinase with similarity to the PRK/URK/PANK kinase subfamily, the PRK/URK/PANK subfamily of P-loop kinases [160]. |
| 12 | XI/382677 [GT to G] | |
| 13 | XII/292021 [TTTGAAAAAAAAAAAAAAAAT to CTTGAAAAAAAAAAAAAAAAAA] | |
| | **Glu3a** | |
| | **Mutation.** | **Gene function. Gene function and adaptation. Gene function and mating behavior.** |
| 1 | IV/383572 [(synonymous_variant c.510G>T p.Val170Val) in PRM7] | Pheromone-regulated protein, role in plasma membrane fusion [161]. |
| 2 | VII/26030 [(missense_variant c.1455G>C p.Met485Ile) in RTG2] | ReTroGrade regulation, Sensor of mitochondrial dysfunction; regulates the subcellular location of Rtg1p and Rtg3p, transcriptional activators of the retrograde (RTG) and TOR pathways [162-164]. |
| 3 | X/125510 [(synonymous_variant c.819C>A p.Gly273Gly) in FAR1] | Factor Arrest, CDK inhibitor and nuclear anchor; during the cell cycle Far1p sequesters the GEF Cdc24p in the nucleus; phosphorylation by Cdc28p-Cln results in SCFCdc4 complex-mediated ubiquitin-dependent degradation [165-168]. Causes activation of GTPase Cdc42p; in response to pheromone, phosphorylation of Far1p by MAPK Fus3p results in association with, and inhibition of Cdc28p-Cln, as well as Msn5p mediated nuclear export of Far1p-Cdc24p, targeting Cdc24p to polarity sites [169]. |
| 4 | X/268831 [(missense_variant c.33A>C p.Leu11Phe) in ARG3] | ARGinine requiring, Ornithine carbamoyltransferase catalyzes the biosynthesis of the arginine precursor citrulline [170,171] |
| 5 | X/669006 [(frameshift_variant c.1369dupG p.Ala457fs) in MNS1] | Alpha-1,2-mannosidase, involved in ER-associated protein degradation (ERAD), catalyzes the removal of one mannose residue from a glycosylated protein, converting the modification from Man9GlcNAc to Man8GlcNAc and catalyzes the last step in glycoprotein maturation in the ER and is critical for ER protein degradation [101,102,104]. Null mutants exhibit an increased lifespan in yeast, Drosophila, and *C. elegans* [172,173]. Increased peptide/protein accumulation in null mutants [101,174]. |

| | | |
|---|---|---|
| 6 | XII/375302 [(missense_variant c.1937G>A p.Gly646Asp) in AVL9] | Apl2 Vps1 Lethal, involved in exocytic transport from the Golgi [44,175]. |
| 7 | IV/300038 [G to GAA 34 bp upstream of ASM4] | Anti-Suppressor in Multicopy, FG-nucleoporin component of central core of nuclear pore complex (NPC), contributes directly to nucleocytoplasmic transport, induces membrane tubulation, which may contribute to nuclear pore assembly [176,177]. Overexpression causes delays in cell cycle progression and abnormal budding pattern [17]. |
| 8 | IV/955986 [GAAA to G 27 bp upstream of VHS1] | Viable in a Hal3 Sit4 background, Cytoplasmic serine/threonine protein kinase, identified as a high-copy suppressor of the synthetic lethality of a sis2 sit4 double mutant, suggesting a role in G1/S phase progression [178]. Sip5 is inhibited by phosphorylation by Vhs1 kinase, whose activity toward Sip5 is stimulated by glucose. Vhs1 might be regulated directly by glucose or by a glucose derivative. Together, these proteins comprise a signaling pathway that regulates Snf1 function [179,180]. Protein serine kinase involved in signal transduction, contributes to G1/S transition of mitotic cell cycle [178]. |
| 9 | VIII/170619 [T to TAAA 279 bp upstream of SLT2] | Suppressor of the LyTic phenotype, Serine/threonine MAP kinase,involved in regulating maintenance of cell wall integrity, cell cycle progression, regulated by the PKC1-mediated signaling pathway [181-183]. MAP kinase localized to the cytoplasm and sites of cell wall growth (bud tips, bud necks, mating projections), Mating response decreased in null mutants (>50% reduction in maximum responsiveness to pheromone) [184]. |
| 10 | II/646091 [CATATATACATATATACATACAT to C] | |
| 11 | X/745451 [G to A] | |
| 12 | XII/1071652 [G to GGTGTGGTGT] | |
| 13 | XIII/538627 [ACACACTCACTCACACATG to A] | |
| **Glu1α** | | |
| | **Mutation.** | **Gene function. Gene function and adaptation. Gene function and mating behavior.** |
| 1 | III/266251 [(missense_variant c.1184A>C p.Lys395Thr) in ABP1] | Actin Binding Protein, Actin-binding protein of the cortical actin cytoskeleton, mediated by Cdc28p and Pho85p [185]. Abp1 reports structural cytoskeleton modifications promoted by glucose withdrawal [186]. Null mutants show decreased sporulation efficiency [187]. |
| 2 | IV/205672 (disruptive_inframe_insertion c.4869_4889 dup GCCAAGCTACA GCCCTACGTC p.Ser1630_Pro1631ins ProSerTyrSerProThrSer) in RPO21 | RNA polymerase II largest subunit B220, reduction of function leads to slow growth, lower mRNA production and decreasing cell viability [188,189]. |
| 3 | VIII/93136 [(missense_variant c.1365_1375del AAAAAAGGAGG ins GAAGAAAGAAA p.Asp459Asn) in YHL008C] | May be involved in the uptake of chloride ions [190-192]. |
| 4 | VIII/93417 [(missense_variant c.1094A>C p.Gln365Pro) in YHL008C] | |
| 5 | VIII/93422 [(synonymous_variant c.1086_1089delACCCinsGCCT p.364) in YHL008C] | |
| 6 | XI/63999 [(missense_variant c.3466_3468delCAGinsAAT p.Gln1156Asn) and (missense_variant c.3461A>T p.Lys1154Met)in MNN4] | Putative positive regulator of mannosylphosphate transferase Mnn6p [193]. |
| 7 | XI/391282 [(stop_gained c.998C>A p.Ser333*) in PAN3] | Poly(A) Nuclease, Essential subunit of the Pan2p-Pan3p poly(A)-ribonuclease complex, controls poly(A) tail length and regulates the stoichiometry and activity of postreplication repair complexes [194-197]. Sporulation efficiency increased in null mutants [9]. |
| 8 | XV/42371 [(missense_variant c.1624T>G p.Ser542Ala) in FRE7] | Putative ferric reductase |
| 9 | VIII/93151 [(missense_variant c.1358_1360delAGAinsGGG p.LysAsn453ArgAsp) in YHL008C] | |
| 10 | IV/376586 [T to TTTC 106 bp upstream of PRP11] | Pre-mRNA Processing, Subunit of the SF3a splicing factor complex, required for spliceosome assembly [198,199]. |

| | | |
|---|---|---|
| 11 | VII/987908 [A to AATATATA TATGTATGCAT ATATATAT 141 bp upstream of MGA1] | Protein similar to heat shock transcription factor [188,200]. |
| 12 | X/161835 [T to TA 79 bp upstream of YJL132W] | |
| 13 | VIII/556927 [intergenic_region n.556927T>G] | |
| 14 | X/204259 [GTAGAAA to G intergenic_region n.204260_204265delTAGAAA] | |
| 15 | XI/663324 [T to A] | |
| 16 | XII/784218 [CGATTC to AAATTA] | |
| 17 | XIV/7057 [intergenic_region n.7057T>C] | |
| 18 | XIV/281738 [C to T] | |
| | **Glu2α** | |
| | **Mutation.** | **Gene function. Gene function and adaptation. Gene function and mating behavior.** |
| 1 | XI/170299 [(synonymous_variant c.831C>G p.Pro277Pro) in SDH1] | Succinate DeHydrogenase [201], couples the oxidation of succinate to the transfer of electrons to ubiquinone as part of the TCA cycle and the mitochondrial respiratory chain [202]. Utilization of carbon source absent in null mutants [203]. |
| 2 | XII/89637 [(stop_gained c.1015G>T p.Glu339*) in HSP104] | Heat Shock Protein required for stress tolerance, and protein disaggregation [204-206]. |
| 3 | III/27658 [TAA to T 299 bp upstream of PEX34, 271 bp upstream of KAR4] | Transcription factor required for response to pheromones, also required during meiosis, exists in two forms, a slower-migrating form more abundant during vegetative growth and a faster-migrating form induced by pheromone [207,208]. Snf1 promotes spindle orientation acting in parallel with Dyn1 and in concert with Kar9. Kar4 presumably regulate cognate genes under a condition unrelated to glucose level [209]. In a null mutant, nuclear fusion during mating: decreased [210], sporulation decreased [9] a karyogamy-specific component of the yeast pheromone response pathway [210]. |
| 4 | VIII/36227 [GA to G 202 upstream of RPL8A] | Ribosomal Protein of the Large subunit, Ribosomal 60S subunit protein L8A, required for processing of 27SA3 pre-rRNA to 27SB pre-rRNA during assembly of large ribosomal subunit; depletion leads to a turnover of pre-rRNA [211]. Snf1/AMPK is activated in the presence of low glucose or alternative carbon sources, thus promoting an energy saving program through transcriptional activation and phosphorylation of metabolic enzymes.interactor of Snf1 [212], null mutant show increased duration in cell cycle progression in G1 phase [213]. |
| 5 | VIII/389016 [AA to TG 21 bp upstream of SUF8] | Proline tRNA (tRNA-Pro) [214]. |
| 6 | XV/894338 [A to AT 246 bp upstream of SLY41] | Suppressor of Loss of Ypt1, Protein involved in ER-to-Golgi transport, packaged into COPII vesicles for trafficking between ER and Golgi [215,216]. |
| 7 | II/38757 [T to TA] | |
| 8 | III/303 [CACACACCCACACCCACACACACCACACCCACACCACACC to ACACCCACACACACAC CACACCCA] | |
| 9 | III/29511 [A to C] | |
| 10 | V/196585 [TAAAAAAAGAAACTGGAAAAAAGGTA to T] | |
| 11 | VIII/21 [A to AC] | |
| 12 | VIII/59082 [AT to A] | |
| 13 | VIII/391898 [A to ATAGTAGTAG] | |
| 14 | XI/664250 [T to A] | |
| 15 | XI/666117 [A to G] | |
| 16 | XII/1064788 [G to A] | |
| 17 | XIV/15990 [TGC to CCC] | |
| 18 | XII/830629 [A to ATTTT] | |
| 19 | VII/434709 [T to CA] | |
| | **Glu3α** | |
| | **Mutation.** | **Gene function. Gene function and adaptation. Gene function and mating behavior.** |
| 1 | X/46206 [(missense_variant c.1228C>A p.Gln410Lys) in LAA1] | Large AP-1 Accessory, AP-1 accessory protein, colocalizes with clathrin to the late-Golgi apparatus also involved in TGN-endosome transport [192,217]. Competitive fitness decreased in null mutants [4]. |

| | | |
|---|---|---|
| 2 | X/667987 [(stop_gained c.344G>A p.Trp115*) in MNS1] | Alpha-1,2-mannosidase, involved in ER-associated protein degradation (ERAD), catalyzes the removal of one mannose residue from a glycosylated protein, converting the modification from Man9GlcNAc to Man8GlcNAc and catalyzes the last step in glycoprotein maturation in the ER and is critical for ER protein degradation [101,102,104]. Increased peptide/protein accumulation in null mutants [101,174]. |
| 3 | XIII/48173 [(missense_variant c.1232G>A p.Cys411Tyr) in BUL2] | Binds Ubiquitin Ligase, Alpha-arrestin, component of the Rsp5p E3-ubiquitin ligase complex, ubiquitin-binding adaptor involved in intracellular amino acid permease sorting [218]. Null mutants show decreased competitive fitness and increased chronological lifespan [4,219]. Null mutants show abnormally elongated buds [220]. |
| 4 | IX/49507 [(disruptive_inframe_insertion c.1439_1441dupAGC p.Gln480dup) in UBP7] | UBiquitin-specific Protease, Ubiquitin-specific protease that cleaves ubiquitin-protein fusions, involved in cell cycle progression through S phase [221,222]. Competitive fitness decreased in null mutants [4]. |
| 5 | XII/823465 [363bp upstream of KAP95] | Karyopherin beta, forms a complex with Srp1p/Kap60p, interacts with nucleoporins to mediate nuclear import of NLS-containing cargo proteins via the nuclear pore complex, regulates PC biosynthesis, GDP-to-GTP exchange factor for Gsp1p [223,224]. |
| 6 | XV/1031350 [356 bp upstream of MRS6] | Mitochondrial RNA Splicing, Rab escort protein, type II geranylgeranyltransferase complex (Bet2p-Bet4p) chaperone, complexes with newly synthesized Rab GTPases, like Ypt1p and Sec4p, modulates the TOR pathway through interactions with Sfp1, alters the kinetics of MAPK pathway activation and polarity reorganization during filamentous growth[225-228] |
| 7 | I/198648 [CGCAA to TGCAG] | |
| 8 | III/224793 [A to GTGTGTGTG] | |
| 9 | XVI/695617 [AAAA to T] | |
| 10 | XVI/775873 [T to AATATCTCA] | |

Upstream in the above tables refers to distance from the start codon of the gene.

**Supplement References.**


1. Hodgson, J. A., Berry, D. R. & Johnston, J. R. Discrimination by heat and proteinase treatments between flocculent phenotypes conferred on Saccharomyces cerevisiae by the genes FLO1 and FLO5. *J Gen Microbiol* **131**, 3219-3227, doi:10.1099/00221287-131-12-3219 (1985).
2. Kobayashi, O., Hayashi, N., Kuroki, R. & Sone, H. Region of FLO1 proteins responsible for sugar recognition. *J Bacteriol* **180**, 6503-6510, doi:10.1128/JB.180.24.6503-6510.1998 (1998).
3. Stratford, M. Evidence for two mechanisms of flocculation in Saccharomyces cerevisiae. *Yeast* **5 Spec No**, S441-445 (1989).
4. Breslow, D. K. *et al.* A comprehensive strategy enabling high-resolution functional analysis of the yeast genome. *Nat Methods* **5**, 711-718, doi:10.1038/nmeth.1234 (2008).
5. Goossens, K. V. *et al.* Molecular mechanism of flocculation self-recognition in yeast and its role in mating and survival. *mBio* **6**, doi:10.1128/mBio.00427-15 (2015).
6. Scholes, D. T., Banerjee, M., Bowen, B. & Curcio, M. J. Multiple regulators of Ty1 transposition in Saccharomyces cerevisiae have conserved roles in genome maintenance. *Genetics* **159**, 1449-1465 (2001).
7. Li, S. *et al.* Rtt105 functions as a chaperone for replication protein A to preserve genome stability. *EMBO J* **37**, doi:10.15252/embj.201899154 (2018).
8. Nakatsukasa, K., Nishikawa, S., Hosokawa, N., Nagata, K. & Endo, T. Mnl1p, an alpha -mannosidase-like protein in yeast Saccharomyces cerevisiae, is required for endoplasmic reticulum-associated degradation of glycoproteins. *J Biol Chem* **276**, 8635-8638, doi:10.1074/jbc.C100023200 (2001).
9. Deutschbauer, A. M., Williams, R. M., Chu, A. M. & Davis, R. W. Parallel phenotypic analysis of sporulation and postgermination growth in Saccharomyces cerevisiae. *Proc Natl Acad Sci U S A* **99**, 15530-15535, doi:10.1073/pnas.202604399 (2002).
10. Fu, D., Beeler, T. & Dunn, T. Sequence, mapping and disruption of CCC1, a gene that cross-complements the Ca(2+)-sensitive phenotype of csg1 mutants. *Yeast* **10**, 515-521, doi:10.1002/yea.320100411 (1994).
11. Lapinskas, P. J., Lin, S. J. & Culotta, V. C. The role of the Saccharomyces cerevisiae CCC1 gene in the homeostasis of manganese ions. *Mol Microbiol* **21**, 519-528, doi:10.1111/j.1365-2958.1996.tb02561.x (1996).
12. Chen, O. S. & Kaplan, J. CCC1 suppresses mitochondrial damage in the yeast model of Friedreich's ataxia by limiting mitochondrial iron accumulation. *J Biol Chem* **275**, 7626-7632, doi:10.1074/jbc.275.11.7626 (2000).
13. Li, L., Chen, O. S., McVey Ward, D. & Kaplan, J. CCC1 is a transporter that mediates vacuolar iron storage in yeast. *J Biol Chem* **276**, 29515-29519, doi:10.1074/jbc.M103944200 (2001).
14. Yoo, C. J. & Wolin, S. L. La proteins from Drosophila melanogaster and Saccharomyces cerevisiae: a yeast homolog of the La autoantigen is dispensable for growth. *Mol Cell Biol* **14**, 5412-5424, doi:10.1128/mcb.14.8.5412-5424.1994 (1994).
15. Yoo, C. J. & Wolin, S. L. The yeast La protein is required for the 3' endonucleolytic cleavage that matures tRNA precursors. *Cell* **89**, 393-402, doi:10.1016/s0092-8674(00)80220-2 (1997).
16. Pannone, B. K., Xue, D. & Wolin, S. L. A role for the yeast La protein in U6 snRNP assembly: evidence that the La protein is a molecular chaperone for RNA polymerase III transcripts. *EMBO J* **17**, 7442-7453, doi:10.1093/emboj/17.24.7442 (1998).
17. Sopko, R. *et al.* Mapping pathways and phenotypes by systematic gene overexpression. *Mol Cell* **21**, 319-330, doi:10.1016/j.molcel.2005.12.011 (2006).

18 Yoshikawa, K. *et al.* Comprehensive phenotypic analysis of single-gene deletion and overexpression strains of Saccharomyces cerevisiae. *Yeast* **28**, 349-361, doi:10.1002/yea.1843 (2011).

19 Sullivan, D. P., Georgiev, A. & Menon, A. K. Tritium suicide selection identifies proteins involved in the uptake and intracellular transport of sterols in Saccharomyces cerevisiae. *Eukaryot Cell* **8**, 161-169, doi:10.1128/EC.00135-08 (2009).

20 Bishop, A. L., Rab, F. A., Sumner, E. R. & Avery, S. V. Phenotypic heterogeneity can enhance rare-cell survival in 'stress-sensitive' yeast populations. *Mol Microbiol* **63**, 507-520, doi:10.1111/j.1365-2958.2006.05504.x (2007).

21 Ho, C. K., Lam, A. F. & Symington, L. S. Identification of nucleases and phosphatases by direct biochemical screen of the Saccharomyces cerevisiae proteome. *PLoS One* **4**, e6993, doi:10.1371/journal.pone.0006993 (2009).

22 Hess, S. M., Stanford, D. R. & Hopper, A. K. SRD1, a S. cerevisiae gene affecting pre-rRNA processing contains a C2/C2 zinc finger motif. *Nucleic Acids Res* **22**, 1265-1271, doi:10.1093/nar/22.7.1265 (1994).

23 Bonilla, M., Nastase, K. K. & Cunningham, K. W. Essential role of calcineurin in response to endoplasmic reticulum stress. *EMBO J* **21**, 2343-2353, doi:10.1093/emboj/21.10.2343 (2002).

24 Viladevall, L. *et al.* Characterization of the calcium-mediated response to alkaline stress in Saccharomyces cerevisiae. *J Biol Chem* **279**, 43614-43624, doi:10.1074/jbc.M403606200 (2004).

25 Cole, J. T., Kean, W. S., Pollard, H. B., Verma, A. & Watson, W. D. Glucose-6-phosphate reduces calcium accumulation in rat brain endoplasmic reticulum. *Front Mol Neurosci* **5**, 51, doi:10.3389/fnmol.2012.00051 (2012).

26 Wolf, B. A., Colca, J. R., Comens, P. G., Turk, J. & McDaniel, M. L. Glucose 6-phosphate regulates Ca2+ steady state in endoplasmic reticulum of islets. A possible link in glucose-induced insulin secretion. *J Biol Chem* **261**, 16284-16287 (1986).

27 Paidhungat, M. & Garrett, S. A homolog of mammalian, voltage-gated calcium channels mediates yeast pheromone-stimulated Ca2+ uptake and exacerbates the cdc1(Ts) growth defect. *Mol Cell Biol* **17**, 6339-6347, doi:10.1128/MCB.17.11.6339 (1997).

28 Fischer, M. *et al.* The Saccharomyces cerevisiae CCH1 gene is involved in calcium influx and mating. *FEBS Lett* **419**, 259-262, doi:10.1016/s0014-5793(97)01466-x (1997).

29 Webber, A. L., Lambrechts, M. G. & Pretorius, I. S. MSS11, a novel yeast gene involved in the regulation of starch metabolism. *Curr Genet* **32**, 260-266, doi:10.1007/s002940050275 (1997).

30 Gagiano, M., van Dyk, D., Bauer, F. F., Lambrechts, M. G. & Pretorius, I. S. Msn1p/Mss10p, Mss11p and Muc1p/Flo11p are part of a signal transduction pathway downstream of Mep2p regulating invasive growth and pseudohyphal differentiation in Saccharomyces cerevisiae. *Mol Microbiol* **31**, 103-116, doi:10.1046/j.1365-2958.1999.01151.x (1999).

31 Gagiano, M. *et al.* Mss11p is a transcription factor regulating pseudohyphal differentiation, invasive growth and starch metabolism in Saccharomyces cerevisiae in response to nutrient availability. *Mol Microbiol* **47**, 119-134, doi:10.1046/j.1365-2958.2003.03247.x (2003).

32 Kim, H. Y., Lee, S. B., Kang, H. S., Oh, G. T. & Kim, T. Two distinct domains of Flo8 activator mediates its role in transcriptional activation and the physical interaction with Mss11. *Biochem Biophys Res Commun* **449**, 202-207, doi:10.1016/j.bbrc.2014.04.161 (2014).

33 Daugeron, M. C., Mauxion, F. & Seraphin, B. The yeast POP2 gene encodes a nuclease involved in mRNA deadenylation. *Nucleic Acids Res* **29**, 2448-2455, doi:10.1093/nar/29.12.2448 (2001).

34 Tucker, M., Staples, R. R., Valencia-Sanchez, M. A., Muhlrad, D. & Parker, R. Ccr4p is the catalytic subunit of a Ccr4p/Pop2p/Notp mRNA deadenylase complex in Saccharomyces cerevisiae. *EMBO J* **21**, 1427-1436, doi:10.1093/emboj/21.6.1427 (2002).

35 Sakai, A., Chibazakura, T., Shimizu, Y. & Hishinuma, F. Molecular analysis of POP2 gene, a gene required for glucose-derepression of gene expression in Saccharomyces cerevisiae. *Nucleic Acids Res* **20**, 6227-6233, doi:10.1093/nar/20.23.6227 (1992).

36 Enyenihi, A. H. & Saunders, W. S. Large-scale functional genomic analysis of sporulation and meiosis in Saccharomyces cerevisiae. *Genetics* **163**, 47-54 (2003).

37 Kloimwieder, A. & Winston, F. A Screen for Germination Mutants in Saccharomyces cerevisiae. *G3 (Bethesda)* **1**, 143-149, doi:10.1534/g3.111.000323 (2011).

38 Buchhaupt, M., Kotter, P. & Entian, K. D. Mutations in the nucleolar proteins Tma23 and Nop6 suppress the malfunction of the Nep1 protein. *FEMS Yeast Res* **7**, 771-781, doi:10.1111/j.1567-1364.2007.00230.x (2007).

39 Fleischer, T. C., Weaver, C. M., McAfee, K. J., Jennings, J. L. & Link, A. J. Systematic identification and functional screens of uncharacterized proteins associated with eukaryotic ribosomal complexes. *Genes Dev* **20**, 1294-1307, doi:10.1101/gad.1422006 (2006).

40 Burtner, C. R., Murakami, C. J., Olsen, B., Kennedy, B. K. & Kaeberlein, M. A genomic analysis of chronological longevity factors in budding yeast. *Cell Cycle* **10**, 1385-1396, doi:10.4161/cc.10.9.15464 (2011).

41 Luke, M. M. *et al.* The SAP, a new family of proteins, associate and function positively with the SIT4 phosphatase. *Mol Cell Biol* **16**, 2744-2755, doi:10.1128/MCB.16.6.2744 (1996).

42 Desfougeres, Y., Gerasimaite, R. U., Jessen, H. J. & Mayer, A. Vtc5, a Novel Subunit of the Vacuolar Transporter Chaperone Complex, Regulates Polyphosphate Synthesis and Phosphate Homeostasis in Yeast. *J Biol Chem* **291**, 22262-22275, doi:10.1074/jbc.M116.746784 (2016).

43 Cohen, A., Perzov, N., Nelson, H. & Nelson, N. A novel family of yeast chaperons involved in the distribution of V-ATPase and other membrane proteins. *J Biol Chem* **274**, 26885-26893, doi:10.1074/jbc.274.38.26885 (1999).

44 Tkach, J. M. *et al.* Dissecting DNA damage response pathways by analysing protein localization and abundance changes during DNA replication stress. *Nat Cell Biol* **14**, 966-976, doi:10.1038/ncb2549 (2012).

45 Buscemi, G., Saracino, F., Masnada, D. & Carbone, M. L. The Saccharomyces cerevisiae SDA1 gene is required for actin cytoskeleton organization and cell cycle progression. *J Cell Sci* **113 ( Pt 7)**, 1199-1211 (2000).

46 Zimmerman, Z. A. & Kellogg, D. R. The Sda1 protein is required for passage through start. *Mol Biol Cell* **12**, 201-219, doi:10.1091/mbc.12.1.201 (2001).

47 Babbio, F., Farinacci, M., Saracino, F., Carbone, M. L. & Privitera, E. Expression and localization studies of hSDA, the human ortholog of the yeast SDA1 gene. *Cell Cycle* **3**, 486-490 (2004).

48 Winston, F. *et al.* Three genes are required for trans-activation of Ty transcription in yeast. *Genetics* **115**, 649-656 (1987).

49 Eisenmann, D. M., Chapon, C., Roberts, S. M., Dollard, C. & Winston, F. The Saccharomyces cerevisiae SPT8 gene encodes a very acidic protein that is functionally related to SPT3 and TATA-binding protein. *Genetics* **137**, 647-657 (1994).

50 Burri, L. & Lithgow, T. A complete set of SNAREs in yeast. *Traffic* **5**, 45-52, doi:10.1046/j.1600-0854.2003.00151.x (2004).

51 Heath, V. L., Shaw, S. L., Roy, S. & Cyert, M. S. Hph1p and Hph2p, novel components of calcineurin-mediated stress responses in Saccharomyces cerevisiae. *Eukaryot Cell* **3**, 695-704, doi:10.1128/EC.3.3.695-704.2004 (2004).

52 Pina, F. J. *et al.* Hph1 and Hph2 are novel components of the Sec63/Sec62 posttranslational translocation complex that aid in vacuolar proton ATPase biogenesis. *Eukaryot Cell* **10**, 63-71, doi:10.1128/EC.00241-10 (2011).

53 Byrne, K. P. & Wolfe, K. H. The Yeast Gene Order Browser: combining curated homology and syntenic context reveals gene fate in polyploid species. *Genome Res* **15**, 1456-1461, doi:10.1101/gr.3672305 (2005).

54 Milkereit, P. *et al.* Maturation and intranuclear transport of pre-ribosomes requires Noc proteins. *Cell* **105**, 499-509, doi:10.1016/s0092-8674(01)00358-0 (2001).

55 Milkereit, P. *et al.* A Noc complex specifically involved in the formation and nuclear export of ribosomal 40 S subunits. *J Biol Chem* **278**, 4072-4081, doi:10.1074/jbc.M208898200 (2003).

56 Dastidar, R. G. *et al.* The nuclear localization of SWI/SNF proteins is subjected to oxygen regulation. *Cell Biosci* **2**, 30, doi:10.1186/2045-3701-2-30 (2012).

57 Nozawa, A., Takano, J., Kobayashi, M., von Wiren, N. & Fujiwara, T. Roles of BOR1, DUR3, and FPS1 in boron transport and tolerance in Saccharomyces cerevisiae. *FEMS Microbiol Lett* **262**, 216-222, doi:10.1111/j.1574-6968.2006.00395.x (2006).

58 VanderSluis, B. *et al.* Broad metabolic sensitivity profiling of a prototrophic yeast deletion collection. *Genome Biol* **15**, R64, doi:10.1186/gb-2014-15-4-r64 (2014).

59 Meyer, S. L., Kvalnes-Krick, K. L. & Schramm, V. L. Characterization of AMD, the AMP deaminase gene in yeast. Production of amd strain, cloning, nucleotide sequence, and properties of the protein. *Biochemistry* **28**, 8734-8743, doi:10.1021/bi00448a009 (1989).

60 Planta, R. J. & Mager, W. H. The list of cytoplasmic ribosomal proteins of Saccharomyces cerevisiae. *Yeast* **14**, 471-477, doi:10.1002/(SICI)1097-0061(19980330)14:5<471::AID-YEA241>3.0.CO;2-U (1998).

61 Ghislain, M., Talla, E. & Francois, J. M. Identification and functional analysis of the Saccharomyces cerevisiae nicotinamidase gene, PNC1. *Yeast* **19**, 215-224, doi:10.1002/yea.810 (2002).

62 Anderson, R. M., Bitterman, K. J., Wood, J. G., Medvedik, O. & Sinclair, D. A. Nicotinamide and PNC1 govern lifespan extension by calorie restriction in Saccharomyces cerevisiae. *Nature* **423**, 181-185, doi:10.1038/nature01578 (2003).

63 Davis, D. A., Bruno, V. M., Loza, L., Filler, S. G. & Mitchell, A. P. Candida albicans Mds3p, a conserved regulator of pH responses and virulence identified through insertional mutagenesis. *Genetics* **162**, 1573-1581 (2002).

64 Benni, M. L. & Neigeborn, L. Identification of a new class of negative regulators affecting sporulation-specific gene expression in yeast. *Genetics* **147**, 1351-1366 (1997).

65 Zhao, H. & Eide, D. J. Zap1p, a metalloregulatory protein involved in zinc-responsive transcriptional regulation in Saccharomyces cerevisiae. *Mol Cell Biol* **17**, 5044-5052, doi:10.1128/MCB.17.9.5044 (1997).

66 Zhao, H. *et al.* Regulation of zinc homeostasis in yeast by binding of the ZAP1 transcriptional activator to zinc-responsive promoter elements. *J Biol Chem* **273**, 28713-28720, doi:10.1074/jbc.273.44.28713 (1998).

67 Nickas, M. E. & Yaffe, M. P. BRO1, a novel gene that interacts with components of the Pkc1p-mitogen-activated protein kinase pathway in Saccharomyces cerevisiae. *Mol Cell Biol* **16**, 2585-2593, doi:10.1128/MCB.16.6.2585 (1996).
68 Odorizzi, G., Katzmann, D. J., Babst, M., Audhya, A. & Emr, S. D. Bro1 is an endosome-associated protein that functions in the MVB pathway in Saccharomyces cerevisiae. *J Cell Sci* **116**, 1893-1903, doi:10.1242/jcs.00395 (2003).
69 Luhtala, N. & Odorizzi, G. Bro1 coordinates deubiquitination in the multivesicular body pathway by recruiting Doa4 to endosomes. *J Cell Biol* **166**, 717-729, doi:10.1083/jcb.200403139 (2004).
70 Pashkova, N. *et al.* The yeast Alix homolog Bro1 functions as a ubiquitin receptor for protein sorting into multivesicular endosomes. *Dev Cell* **25**, 520-533, doi:10.1016/j.devcel.2013.04.007 (2013).
71 Houseley, J., Rubbi, L., Grunstein, M., Tollervey, D. & Vogelauer, M. A ncRNA modulates histone modification and mRNA induction in the yeast GAL gene cluster. *Mol Cell* **32**, 685-695, doi:10.1016/j.molcel.2008.09.027 (2008).
72 Pinskaya, M., Gourvennec, S. & Morillon, A. H3 lysine 4 di- and tri-methylation deposited by cryptic transcription attenuates promoter activation. *EMBO J* **28**, 1697-1707, doi:10.1038/emboj.2009.108 (2009).
73 Strom, M., Vollmer, P., Tan, T. J. & Gallwitz, D. A yeast GTPase-activating protein that interacts specifically with a member of the Ypt/Rab family. *Nature* **361**, 736-739, doi:10.1038/361736a0 (1993).
74 Will, E. & Gallwitz, D. Biochemical characterization of Gyp6p, a Ypt/Rab-specific GTPase-activating protein from yeast. *J Biol Chem* **276**, 12135-12139, doi:10.1074/jbc.M011451200 (2001).
75 Clavilier, L., Pere-Aubert, G., Somlo, M. & Slonimski, P. P. [Network of interactions between unlinked genes: synergistic and antagonistic regulation of iso-1-cytochrome c, iso-2-cytochrome c and cytochrome b2 synthesis]. *Biochimie* **58**, 155-172, doi:10.1016/s0300-9084(76)80366-5 (1976).
76 Dumont, M. E., Cardillo, T. S., Hayes, M. K. & Sherman, F. Role of cytochrome c heme lyase in mitochondrial import and accumulation of cytochrome c in Saccharomyces cerevisiae. *Mol Cell Biol* **11**, 5487-5496, doi:10.1128/mcb.11.11.5487-5496.1991 (1991).
77 Moir, D., Stewart, S. E., Osmond, B. C. & Botstein, D. Cold-sensitive cell-division-cycle mutants of yeast: isolation, properties, and pseudoreversion studies. *Genetics* **100**, 547-563 (1982).
78 Bruck, I. & Kaplan, D. L. Cdc45 protein-single-stranded DNA interaction is important for stalling the helicase during replication stress. *J Biol Chem* **288**, 7550-7563, doi:10.1074/jbc.M112.440941 (2013).
79 Belogrudov, G. I. *et al.* Yeast COQ4 encodes a mitochondrial protein required for coenzyme Q synthesis. *Arch Biochem Biophys* **392**, 48-58, doi:10.1006/abbi.2001.2448 (2001).
80 Marbois, B. *et al.* Coq3 and Coq4 define a polypeptide complex in yeast mitochondria for the biosynthesis of coenzyme Q. *J Biol Chem* **280**, 20231-20238, doi:10.1074/jbc.M501315200 (2005).
81 Casarin, A. *et al.* Functional characterization of human COQ4, a gene required for Coenzyme Q10 biosynthesis. *Biochem Biophys Res Commun* **372**, 35-39, doi:10.1016/j.bbrc.2008.04.172 (2008).
82 Berenguel Hernandez, A. M. *et al.* Design of High-Throughput Screening of Natural Extracts to Identify Molecules Bypassing Primary Coenzyme Q Deficiency in

Saccharomyces cerevisiae. *SLAS Discov* **25**, 299-309, doi:10.1177/2472555219877185 (2020).

83 Edwards, M. C. *et al.* Human CPR (cell cycle progression restoration) genes impart a Far- phenotype on yeast cells. *Genetics* **147**, 1063-1076 (1997).

84 Wu, C., Jansen, G., Zhang, J., Thomas, D. Y. & Whiteway, M. Adaptor protein Ste50p links the Ste11p MEKK to the HOG pathway through plasma membrane association. *Genes Dev* **20**, 734-746, doi:10.1101/gad.1375706 (2006).

85 Karunanithi, S. & Cullen, P. J. The filamentous growth MAPK Pathway Responds to Glucose Starvation Through the Mig1/2 transcriptional repressors in Saccharomyces cerevisiae. *Genetics* **192**, 869-887, doi:10.1534/genetics.112.142661 (2012).

86 Ohi, M. D. *et al.* Proteomics analysis reveals stable multiprotein complexes in both fission and budding yeasts containing Myb-related Cdc5p/Cef1p, novel pre-mRNA splicing factors, and snRNAs. *Mol Cell Biol* **22**, 2011-2024, doi:10.1128/MCB.22.7.2011-2024.2002 (2002).

87 Haurie, V. *et al.* The transcriptional activator Cat8p provides a major contribution to the reprogramming of carbon metabolism during the diauxic shift in Saccharomyces cerevisiae. *J Biol Chem* **276**, 76-85, doi:10.1074/jbc.M008752200 (2001).

88 Tizon, B., Rodriguez-Torres, A. M. & Cerdan, M. E. Disruption of six novel Saccharomyces cerevisiae genes reveals that YGL129c is necessary for growth in non-fermentable carbon sources, YGL128c for growth at low or high temperatures and YGL125w is implicated in the biosynthesis of methionine. *Yeast* **15**, 145-154, doi:10.1002/(SICI)1097-0061(19990130)15:2<145::AID-YEA346>3.0.CO;2-J (1999).

89 Briza, P. *et al.* Systematic analysis of sporulation phenotypes in 624 non-lethal homozygous deletion strains of Saccharomyces cerevisiae. *Yeast* **19**, 403-422, doi:10.1002/yea.843 (2002).

90 Siniossoglou, S., Hurt, E. C. & Pelham, H. R. Psr1p/Psr2p, two plasma membrane phosphatases with an essential DXDX(T/V) motif required for sodium stress response in yeast. *J Biol Chem* **275**, 19352-19360, doi:10.1074/jbc.M001314200 (2000).

91 Boeckstaens, M., Llinares, E., Van Vooren, P. & Marini, A. M. The TORC1 effector kinase Npr1 fine tunes the inherent activity of the Mep2 ammonium transport protein. *Nat Commun* **5**, 3101, doi:10.1038/ncomms4101 (2014).

92 Chen, X. *et al.* Whi2 is a conserved negative regulator of TORC1 in response to low amino acids. *PLoS Genet* **14**, e1007592, doi:10.1371/journal.pgen.1007592 (2018).

93 Li, R. Bifurcation of the mitotic checkpoint pathway in budding yeast. *Proc Natl Acad Sci U S A* **96**, 4989-4994, doi:10.1073/pnas.96.9.4989 (1999).

94 Pereira, G., Hofken, T., Grindlay, J., Manson, C. & Schiebel, E. The Bub2p spindle checkpoint links nuclear migration with mitotic exit. *Mol Cell* **6**, 1-10 (2000).

95 Wang, Y., Hu, F. & Elledge, S. J. The Bfa1/Bub2 GAP complex comprises a universal checkpoint required to prevent mitotic exit. *Curr Biol* **10**, 1379-1382, doi:10.1016/s0960-9822(00)00779-x (2000).

96 Geymonat, M. *et al.* Control of mitotic exit in budding yeast. In vitro regulation of Tem1 GTPase by Bub2 and Bfa1. *J Biol Chem* **277**, 28439-28445, doi:10.1074/jbc.M202540200 (2002).

97 Fraschini, R., D'Ambrosio, C., Venturetti, M., Lucchini, G. & Piatti, S. Disappearance of the budding yeast Bub2-Bfa1 complex from the mother-bound spindle pole contributes to mitotic exit. *J Cell Biol* **172**, 335-346, doi:10.1083/jcb.200507162 (2006).

98 Mondeel, T., Holland, P., Nielsen, J. & Barberis, M. ChIP-exo analysis highlights Fkh1 and Fkh2 transcription factors as hubs that integrate multi-scale networks in budding yeast. *Nucleic Acids Res* **47**, 7825-7841, doi:10.1093/nar/gkz603 (2019).

99 Attner, M. A. & Amon, A. Control of the mitotic exit network during meiosis. *Mol Biol Cell* **23**, 3122-3132, doi:10.1091/mbc.E12-03-0235 (2012).

100 Gordon, O. *et al.* Nud1p, the yeast homolog of Centriolin, regulates spindle pole body inheritance in meiosis. *EMBO J* **25**, 3856-3868, doi:10.1038/sj.emboj.7601254 (2006).

101 Camirand, A., Heysen, A., Grondin, B. & Herscovics, A. Glycoprotein biosynthesis in Saccharomyces cerevisiae. Isolation and characterization of the gene encoding a specific processing alpha-mannosidase. *J Biol Chem* **266**, 15120-15127 (1991).

102 Burke, J., Lipari, F., Igdoura, S. & Herscovics, A. The Saccharomyces cerevisiae processing alpha 1,2-mannosidase is localized in the endoplasmic reticulum, independently of known retrieval motifs. *Eur J Cell Biol* **70**, 298-305 (1996).

103 Knop, M., Hauser, N. & Wolf, D. H. N-Glycosylation affects endoplasmic reticulum degradation of a mutated derivative of carboxypeptidase yscY in yeast. *Yeast* **12**, 1229-1238, doi:10.1002/(sici)1097-0061(19960930)12:12<1229::aid-yea15>3.0.co;2-h (1996).

104 Jakob, C. A., Burda, P., Roth, J. & Aebi, M. Degradation of misfolded endoplasmic reticulum glycoproteins in Saccharomyces cerevisiae is determined by a specific oligosaccharide structure. *J Cell Biol* **142**, 1223-1233, doi:10.1083/jcb.142.5.1223 (1998).

105 Zuk, D., Belk, J. P. & Jacobson, A. Temperature-sensitive mutations in the Saccharomyces cerevisiae MRT4, GRC5, SLA2 and THS1 genes result in defects in mRNA turnover. *Genetics* **153**, 35-47 (1999).

106 Harnpicharnchai, P. *et al.* Composition and functional characterization of yeast 66S ribosome assembly intermediates. *Mol Cell* **8**, 505-515, doi:10.1016/s1097-2765(01)00344-6 (2001).

107 Oltmanns, O. & Bacher, A. Biosynthesis of riboflavine in Saccharomyces cerevisiae: the role of genes rib 1 and rib 7. *J Bacteriol* **110**, 818-822, doi:10.1128/jb.110.3.818-822.1972 (1972).

108 Buitrago, M. J., Gonzalez, G. A., Saiz, J. E. & Revuelta, J. L. Mapping of the RIB1 and RIB7 genes involved in the biosynthesis of riboflavin in Saccharomyces cerevisiae. *Yeast* **9**, 1099-1102, doi:10.1002/yea.320091009 (1993).

109 Richter, G. *et al.* Biosynthesis of riboflavin: characterization of the bifunctional deaminase-reductase of Escherichia coli and Bacillus subtilis. *J Bacteriol* **179**, 2022-2028, doi:10.1128/jb.179.6.2022-2028.1997 (1997).

110 Paeschke, K. *et al.* Pif1 family helicases suppress genome instability at G-quadruplex motifs. *Nature* **497**, 458-462, doi:10.1038/nature12149 (2013).

111 <https://www.yeastgenome.org/reference/S000134293> (

112 Budovskaya, Y. V., Stephan, J. S., Deminoff, S. J. & Herman, P. K. An evolutionary proteomics approach identifies substrates of the cAMP-dependent protein kinase. *Proc Natl Acad Sci U S A* **102**, 13933-13938, doi:10.1073/pnas.0501046102 (2005).

113 Tomenchok, D. M. & Brandriss, M. C. Gene-enzyme relationships in the proline biosynthetic pathway of Saccharomyces cerevisiae. *J Bacteriol* **169**, 5364-5372, doi:10.1128/jb.169.12.5364-5372.1987 (1987).

114 Chen, S., Tarsio, M., Kane, P. M. & Greenberg, M. L. Cardiolipin mediates cross-talk between mitochondria and the vacuole. *Mol Biol Cell* **19**, 5047-5058, doi:10.1091/mbc.E08-05-0486 (2008).

115 Jiang, F., Gu, Z., Granger, J. M. & Greenberg, M. L. Cardiolipin synthase expression is essential for growth at elevated temperature and is regulated by factors affecting

mitochondrial development. *Mol Microbiol* **31**, 373-379, doi:10.1046/j.1365-2958.1999.01181.x (1999).
116 Jiang, F. *et al.* Absence of cardiolipin in the crd1 null mutant results in decreased mitochondrial membrane potential and reduced mitochondrial function. *J Biol Chem* **275**, 22387-22394, doi:10.1074/jbc.M909868199 (2000).
117 Joshi, A. S., Thompson, M. N., Fei, N., Huttemann, M. & Greenberg, M. L. Cardiolipin and mitochondrial phosphatidylethanolamine have overlapping functions in mitochondrial fusion in Saccharomyces cerevisiae. *J Biol Chem* **287**, 17589-17597, doi:10.1074/jbc.M111.330167 (2012).
118 Gatta, A. T. *et al.* A new family of StART domain proteins at membrane contact sites has a role in ER-PM sterol transport. *Elife* **4**, doi:10.7554/eLife.07253 (2015).
119 Sokolov, S. S., Galkina, K. V., Litvinova, E. A., Knorre, D. A. & Severin, F. F. The Role of LAM Genes in the Pheromone-Induced Cell Death of S. cerevisiae Yeast. *Biochemistry (Mosc)* **85**, 300-309, doi:10.1134/S0006297920030050 (2020).
120 Rodriguez, C. & Flores, C. Mutations in GAL2 or GAL4 alleviate catabolite repression produced by galactose in Saccharomyces cerevisiae. *Enzyme Microb Technol* **26**, 748-755, doi:10.1016/s0141-0229(00)00167-8 (2000).
121 Kasahara, T. & Kasahara, M. Three aromatic amino acid residues critical for galactose transport in yeast Gal2 transporter. *J Biol Chem* **275**, 4422-4428, doi:10.1074/jbc.275.6.4422 (2000).
122 Maier, A., Volker, B., Boles, E. & Fuhrmann, G. F. Characterisation of glucose transport in Saccharomyces cerevisiae with plasma membrane vesicles (countertransport) and intact cells (initial uptake) with single Hxt1, Hxt2, Hxt3, Hxt4, Hxt6, Hxt7 or Gal2 transporters. *FEMS Yeast Res* **2**, 539-550, doi:10.1111/j.1567-1364.2002.tb00121.x (2002).
123 Douglas, H. C. & Condie, F. The genetic control of galactose utilization in Saccharomyces. *J Bacteriol* **68**, 662-670, doi:10.1128/jb.68.6.662-670.1954 (1954).
124 Levin, D. E., Hammond, C. I., Ralston, R. O. & Bishop, J. M. Two yeast genes that encode unusual protein kinases. *Proc Natl Acad Sci U S A* **84**, 6035-6039, doi:10.1073/pnas.84.17.6035 (1987).
125 Elbert, M., Rossi, G. & Brennwald, P. The yeast par-1 homologs kin1 and kin2 show genetic and physical interactions with components of the exocytic machinery. *Mol Biol Cell* **16**, 532-549, doi:10.1091/mbc.e04-07-0549 (2005).
126 Jeschke, G. R. *et al.* Substrate priming enhances phosphorylation by the budding yeast kinases Kin1 and Kin2. *J Biol Chem* **293**, 18353-18364, doi:10.1074/jbc.RA118.005651 (2018).
127 Yuan, S. M., Nie, W. C., He, F., Jia, Z. W. & Gao, X. D. Kin2, the Budding Yeast Ortholog of Animal MARK/PAR-1 Kinases, Localizes to the Sites of Polarized Growth and May Regulate Septin Organization and the Cell Wall. *PLoS One* **11**, e0153992, doi:10.1371/journal.pone.0153992 (2016).
128 Leem, S. H. *et al.* The possible mechanism of action of ciclopirox olamine in the yeast Saccharomyces cerevisiae. *Mol Cells* **15**, 55-61 (2003).
129 Measday, V. *et al.* A family of cyclin-like proteins that interact with the Pho85 cyclin-dependent kinase. *Mol Cell Biol* **17**, 1212-1223, doi:10.1128/MCB.17.3.1212 (1997).
130 Singer-Kruger, B. *et al.* Role of three rab5-like GTPases, Ypt51p, Ypt52p, and Ypt53p, in the endocytic and vacuolar protein sorting pathways of yeast. *J Cell Biol* **125**, 283-298, doi:10.1083/jcb.125.2.283 (1994).
131 Thuriaux, P. *et al.* Gene RPA43 in Saccharomyces cerevisiae encodes an essential subunit of RNA polymerase I. *J Biol Chem* **270**, 24252-24257, doi:10.1074/jbc.270.41.24252 (1995).

132 Protchenko, O. *et al.* Three cell wall mannoproteins facilitate the uptake of iron in Saccharomyces cerevisiae. *J Biol Chem* **276**, 49244-49250, doi:10.1074/jbc.M109220200 (2001).

133 Philpott, C. C., Protchenko, O., Kim, Y. W., Boretsky, Y. & Shakoury-Elizeh, M. The response to iron deprivation in Saccharomyces cerevisiae: expression of siderophore-based systems of iron uptake. *Biochem Soc Trans* **30**, 698-702, doi:10.1042/bst0300698 (2002).

134 Kohno, H. *et al.* Bni1p implicated in cytoskeletal control is a putative target of Rho1p small GTP binding protein in Saccharomyces cerevisiae. *EMBO J* **15**, 6060-6068 (1996).

135 Zahner, J. E., Harkins, H. A. & Pringle, J. R. Genetic analysis of the bipolar pattern of bud site selection in the yeast Saccharomyces cerevisiae. *Mol Cell Biol* **16**, 1857-1870, doi:10.1128/MCB.16.4.1857 (1996).

136 Sagot, I., Rodal, A. A., Moseley, J., Goode, B. L. & Pellman, D. An actin nucleation mechanism mediated by Bni1 and profilin. *Nat Cell Biol* **4**, 626-631, doi:10.1038/ncb834 (2002).

137 Qi, M. & Elion, E. A. Formin-induced actin cables are required for polarized recruitment of the Ste5 scaffold and high level activation of MAPK Fus3. *J Cell Sci* **118**, 2837-2848, doi:10.1242/jcs.02418 (2005).

138 Dorer, R., Boone, C., Kimbrough, T., Kim, J. & Hartwell, L. H. Genetic analysis of default mating behavior in Saccharomyces cerevisiae. *Genetics* **146**, 39-55 (1997).

139 Nelson, B. *et al.* RAM: a conserved signaling network that regulates Ace2p transcriptional activity and polarized morphogenesis. *Mol Biol Cell* **14**, 3782-3803, doi:10.1091/mbc.e03-01-0018 (2003).

140 Ni, L. & Snyder, M. A genomic study of the bipolar bud site selection pattern in Saccharomyces cerevisiae. *Mol Biol Cell* **12**, 2147-2170, doi:10.1091/mbc.12.7.2147 (2001).

141 Hofmann, C. *et al.* Saccharomyces cerevisiae Duo1p and Dam1p, novel proteins involved in mitotic spindle function. *J Cell Biol* **143**, 1029-1040, doi:10.1083/jcb.143.4.1029 (1998).

142 Li, Y. *et al.* The mitotic spindle is required for loading of the DASH complex onto the kinetochore. *Genes Dev* **16**, 183-197, doi:10.1101/gad.959402 (2002).

143 Sattlegger, E. *et al.* Gcn1 and actin binding to Yih1: implications for activation of the eIF2 kinase GCN2. *J Biol Chem* **286**, 10341-10355, doi:10.1074/jbc.M110.171587 (2011).

144 Sattlegger, E. *et al.* YIH1 is an actin-binding protein that inhibits protein kinase GCN2 and impairs general amino acid control when overexpressed. *J Biol Chem* **279**, 29952-29962, doi:10.1074/jbc.M404009200 (2004).

145 Silva, R. C. *et al.* The Gcn2 Regulator Yih1 Interacts with the Cyclin Dependent Kinase Cdc28 and Promotes Cell Cycle Progression through G2/M in Budding Yeast. *PLoS One* **10**, e0131070, doi:10.1371/journal.pone.0131070 (2015).

146 Mai, B. & Breeden, L. Xbp1, a stress-induced transcriptional repressor of the Saccharomyces cerevisiae Swi4/Mbp1 family. *Mol Cell Biol* **17**, 6491-6501, doi:10.1128/MCB.17.11.6491 (1997).

147 Mai, B. & Breeden, L. CLN1 and its repression by Xbp1 are important for efficient sporulation in budding yeast. *Mol Cell Biol* **20**, 478-487, doi:10.1128/MCB.20.2.478-487.2000 (2000).

148 Miles, S., Li, L., Davison, J. & Breeden, L. L. Xbp1 directs global repression of budding yeast transcription during the transition to quiescence and is important for the longevity

and reversibility of the quiescent state. *PLoS Genet* **9**, e1003854, doi:10.1371/journal.pgen.1003854 (2013).
149 Athenstaedt, K. & Daum, G. Tgl4p and Tgl5p, two triacylglycerol lipases of the yeast Saccharomyces cerevisiae are localized to lipid particles. *J Biol Chem* **280**, 37301-37309, doi:10.1074/jbc.M507261200 (2005).
150 Merkler, D. J. & Schramm, V. L. Catalytic and regulatory site composition of yeast AMP deaminase by comparative binding and rate studies. Resolution of the cooperative mechanism. *J Biol Chem* **265**, 4420-4426 (1990).
151 Merkler, D. J., Wali, A. S., Taylor, J. & Schramm, V. L. AMP deaminase from yeast. Role in AMP degradation, large scale purification, and properties of the native and proteolyzed enzyme. *J Biol Chem* **264**, 21422-21430 (1989).
152 Akizu, N. *et al.* AMPD2 regulates GTP synthesis and is mutated in a potentially treatable neurodegenerative brainstem disorder. *Cell* **154**, 505-517, doi:10.1016/j.cell.2013.07.005 (2013).
153 Walther, T. *et al.* Control of ATP homeostasis during the respiro-fermentative transition in yeast. *Mol Syst Biol* **6**, 344, doi:10.1038/msb.2009.100 (2010).
154 Schaaff, I., Hohmann, S. & Zimmermann, F. K. Molecular analysis of the structural gene for yeast transaldolase. *Eur J Biochem* **188**, 597-603, doi:10.1111/j.1432-1033.1990.tb15440.x (1990).
155 Bengtsson, O. *et al.* Identification of common traits in improved xylose-growing Saccharomyces cerevisiae for inverse metabolic engineering. *Yeast* **25**, 835-847, doi:10.1002/yea.1638 (2008).
156 Cai, M. & Davis, R. W. Yeast centromere binding protein CBF1, of the helix-loop-helix protein family, is required for chromosome stability and methionine prototrophy. *Cell* **61**, 437-446, doi:10.1016/0092-8674(90)90525-j (1990).
157 Wieland, G. *et al.* Determination of the binding constants of the centromere protein Cbf1 to all 16 centromere DNAs of Saccharomyces cerevisiae. *Nucleic Acids Res* **29**, 1054-1060, doi:10.1093/nar/29.5.1054 (2001).
158 Moreau, J. L. *et al.* Regulated displacement of TBP from the PHO8 promoter in vivo requires Cbf1 and the Isw1 chromatin remodeling complex. *Mol Cell* **11**, 1609-1620, doi:10.1016/s1097-2765(03)00184-9 (2003).
159 Kent, N. A., Eibert, S. M. & Mellor, J. Cbf1p is required for chromatin remodeling at promoter-proximal CACGTG motifs in yeast. *J Biol Chem* **279**, 27116-27123, doi:10.1074/jbc.M403818200 (2004).
160 Gueguen-Chaignon, V. *et al.* Crystal structure and functional analysis identify the P-loop containing protein YFH7 of Saccharomyces cerevisiae as an ATP-dependent kinase. *Proteins* **71**, 804-812, doi:10.1002/prot.21740 (2008).
161 Heiman, M. G. & Walter, P. Prm1p, a pheromone-regulated multispanning membrane protein, facilitates plasma membrane fusion during yeast mating. *J Cell Biol* **151**, 719-730, doi:10.1083/jcb.151.3.719 (2000).
162 Liao, X. & Butow, R. A. RTG1 and RTG2: two yeast genes required for a novel path of communication from mitochondria to the nucleus. *Cell* **72**, 61-71, doi:10.1016/0092-8674(93)90050-z (1993).
163 Sekito, T., Thornton, J. & Butow, R. A. Mitochondria-to-nuclear signaling is regulated by the subcellular localization of the transcription factors Rtg1p and Rtg3p. *Mol Biol Cell* **11**, 2103-2115, doi:10.1091/mbc.11.6.2103 (2000).
164 Komeili, A., Wedaman, K. P., O'Shea, E. K. & Powers, T. Mechanism of metabolic control. Target of rapamycin signaling links nitrogen quality to the activity of the Rtg1 and Rtg3 transcription factors. *J Cell Biol* **151**, 863-878, doi:10.1083/jcb.151.4.863 (2000).

165 Tyers, M. & Futcher, B. Far1 and Fus3 link the mating pheromone signal transduction pathway to three G1-phase Cdc28 kinase complexes. *Mol Cell Biol* **13**, 5659-5669, doi:10.1128/mcb.13.9.5659-5669.1993 (1993).

166 Nern, A. & Arkowitz, R. A. A Cdc24p-Far1p-Gbetagamma protein complex required for yeast orientation during mating. *J Cell Biol* **144**, 1187-1202, doi:10.1083/jcb.144.6.1187 (1999).

167 Peter, M. & Herskowitz, I. Direct inhibition of the yeast cyclin-dependent kinase Cdc28-Cln by Far1. *Science* **265**, 1228-1231, doi:10.1126/science.8066461 (1994).

168 Chang, F. & Herskowitz, I. Identification of a gene necessary for cell cycle arrest by a negative growth factor of yeast: FAR1 is an inhibitor of a G1 cyclin, CLN2. *Cell* **63**, 999-1011, doi:10.1016/0092-8674(90)90503-7 (1990).

169 Alberghina, L., Rossi, R. L., Querin, L., Wanke, V. & Vanoni, M. A cell sizer network involving Cln3 and Far1 controls entrance into S phase in the mitotic cycle of budding yeast. *J Cell Biol* **167**, 433-443, doi:10.1083/jcb.200405102 (2004).

170 Messenguy, F. Regulation of arginine biosynthesis in Saccharomyces cerevisiae: isolation of a cis-dominant, constitutive mutant for ornithine carbamoyltransferase synthesis. *J Bacteriol* **128**, 49-55, doi:10.1128/jb.128.1.49-55.1976 (1976).

171 Crabeel, M., Messenguy, F., Lacroute, F. & Glansdorff, N. Cloning arg3, the gene for ornithine carbamoyltransferase from Saccharomyces cerevisiae: expression in Escherichia coli requires secondary mutations; production of plasmid beta-lactamase in yeast. *Proc Natl Acad Sci U S A* **78**, 5026-5030, doi:10.1073/pnas.78.8.5026 (1981).

172 Fabrizio, P. *et al.* Genome-wide screen in Saccharomyces cerevisiae identifies vacuolar protein sorting, autophagy, biosynthetic, and tRNA methylation genes involved in life span regulation. *PLoS Genet* **6**, e1001024, doi:10.1371/journal.pgen.1001024 (2010).

173 Liu, Y. L. *et al.* Reduced expression of alpha-1,2-mannosidase I extends lifespan in Drosophila melanogaster and Caenorhabditis elegans. *Aging Cell* **8**, 370-379, doi:10.1111/j.1474-9726.2009.00471.x (2009).

174 Martinez Benitez, E., Stolz, A., Becher, A. & Wolf, D. H. Mnl2, a novel component of the ER associated protein degradation pathway. *Biochem Biophys Res Commun* **414**, 528-532, doi:10.1016/j.bbrc.2011.09.100 (2011).

175 Harsay, E. & Schekman, R. Avl9p, a member of a novel protein superfamily, functions in the late secretory pathway. *Mol Biol Cell* **18**, 1203-1219, doi:10.1091/mbc.e06-11-1035 (2007).

176 Giot, L., Simon, M., Dubois, C. & Faye, G. Suppressors of thermosensitive mutations in the DNA polymerase delta gene of Saccharomyces cerevisiae. *Mol Gen Genet* **246**, 212-222, doi:10.1007/BF00294684 (1995).

177 Marelli, M., Aitchison, J. D. & Wozniak, R. W. Specific binding of the karyopherin Kap121p to a subunit of the nuclear pore complex containing Nup53p, Nup59p, and Nup170p. *J Cell Biol* **143**, 1813-1830, doi:10.1083/jcb.143.7.1813 (1998).

178 Munoz, I., Simon, E., Casals, N., Clotet, J. & Arino, J. Identification of multicopy suppressors of cell cycle arrest at the G1-S transition in Saccharomyces cerevisiae. *Yeast* **20**, 157-169, doi:10.1002/yea.938 (2003).

179 Simpson-Lavy, K. & Kupiec, M. A reversible liquid drop aggregation controls glucose response in yeast. *Curr Genet* **64**, 785-788, doi:10.1007/s00294-018-0805-0 (2018).

180 Simpson-Lavy, K., Xu, T., Johnston, M. & Kupiec, M. The Std1 Activator of the Snf1/AMPK Kinase Controls Glucose Response in Yeast by a Regulated Protein Aggregation. *Mol Cell* **68**, 1120-1133 e1123, doi:10.1016/j.molcel.2017.11.016 (2017).

181 Martin-Yken, H., Dagkessamanskaia, A., Basmaji, F., Lagorce, A. & Francois, J. The interaction of Slt2 MAP kinase with Knr4 is necessary for signalling through the cell

wall integrity pathway in Saccharomyces cerevisiae. *Mol Microbiol* **49**, 23-35, doi:10.1046/j.1365-2958.2003.03541.x (2003).
182 Carmody, S. R., Tran, E. J., Apponi, L. H., Corbett, A. H. & Wente, S. R. The mitogen-activated protein kinase Slt2 regulates nuclear retention of non-heat shock mRNAs during heat shock-induced stress. *Mol Cell Biol* **30**, 5168-5179, doi:10.1128/MCB.00735-10 (2010).
183 Madden, K., Sheu, Y. J., Baetz, K., Andrews, B. & Snyder, M. SBF cell cycle regulator as a target of the yeast PKC-MAP kinase pathway. *Science* **275**, 1781-1784, doi:10.1126/science.275.5307.1781 (1997).
184 Chasse, S. A. *et al.* Genome-scale analysis reveals Sst2 as the principal regulator of mating pheromone signaling in the yeast Saccharomyces cerevisiae. *Eukaryot Cell* **5**, 330-346, doi:10.1128/EC.5.2.330-346.2006 (2006).
185 Drubin, D. G., Miller, K. G. & Botstein, D. Yeast actin-binding proteins: evidence for a role in morphogenesis. *J Cell Biol* **107**, 2551-2561, doi:10.1083/jcb.107.6.2551 (1988).
186 Espinoza-Simon, E. *et al.* In Saccharomyces cerevisiae, withdrawal of the carbon source results in detachment of glycolytic enzymes from the cytoskeleton and in actin reorganization. *Fungal Biol* **124**, 15-23, doi:10.1016/j.funbio.2019.10.005 (2020).
187 Lila, T. & Drubin, D. G. Evidence for physical and functional interactions among two Saccharomyces cerevisiae SH3 domain proteins, an adenylyl cyclase-associated protein and the actin cytoskeleton. *Mol Biol Cell* **8**, 367-385, doi:10.1091/mbc.8.2.367 (1997).
188 Woychik, N. A. & Hampsey, M. The RNA polymerase II machinery: structure illuminates function. *Cell* **108**, 453-463, doi:10.1016/s0092-8674(02)00646-3 (2002).
189 Ingles, C. J., Himmelfarb, H. J., Shales, M., Greenleaf, A. L. & Friesen, J. D. Identification, molecular cloning, and mutagenesis of Saccharomyces cerevisiae RNA polymerase genes. *Proc Natl Acad Sci U S A* **81**, 2157-2161, doi:10.1073/pnas.81.7.2157 (1984).
190 Makuc, J. *et al.* The putative monocarboxylate permeases of the yeast Saccharomyces cerevisiae do not transport monocarboxylic acids across the plasma membrane. *Yeast* **18**, 1131-1143, doi:10.1002/yea.763 (2001).
191 Jennings, M. L. & Cui, J. Chloride homeostasis in Saccharomyces cerevisiae: high affinity influx, V-ATPase-dependent sequestration, and identification of a candidate Cl- sensor. *J Gen Physiol* **131**, 379-391, doi:10.1085/jgp.200709905 (2008).
192 Huh, W. K. *et al.* Global analysis of protein localization in budding yeast. *Nature* **425**, 686-691, doi:10.1038/nature02026 (2003).
193 Raschke, W. C., Kern, K. A., Antalis, C. & Ballou, C. E. Genetic control of yeast mannan structure. Isolation and characterization of mannan mutants. *J Biol Chem* **248**, 4660-4666 (1973).
194 Brown, C. E., Tarun, S. Z., Jr., Boeck, R. & Sachs, A. B. PAN3 encodes a subunit of the Pab1p-dependent poly(A) nuclease in Saccharomyces cerevisiae. *Mol Cell Biol* **16**, 5744-5753, doi:10.1128/MCB.16.10.5744 (1996).
195 Brown, C. E. & Sachs, A. B. Poly(A) tail length control in Saccharomyces cerevisiae occurs by message-specific deadenylation. *Mol Cell Biol* **18**, 6548-6559, doi:10.1128/MCB.18.11.6548 (1998).
196 Hammet, A., Pike, B. L. & Heierhorst, J. Posttranscriptional regulation of the RAD5 DNA repair gene by the Dun1 kinase and the Pan2-Pan3 poly(A)-nuclease complex contributes to survival of replication blocks. *J Biol Chem* **277**, 22469-22474, doi:10.1074/jbc.M202473200 (2002).

197 Wolf, J. *et al.* Structural basis for Pan3 binding to Pan2 and its function in mRNA recruitment and deadenylation. *EMBO J* **33**, 1514-1526, doi:10.15252/embj.201488373 (2014).

198 Vijayraghavan, U., Company, M. & Abelson, J. Isolation and characterization of pre-mRNA splicing mutants of Saccharomyces cerevisiae. *Genes Dev* **3**, 1206-1216, doi:10.1101/gad.3.8.1206 (1989).

199 Hodges, P. E. & Beggs, J. D. RNA splicing. U2 fulfils a commitment. *Curr Biol* **4**, 264-267, doi:10.1016/s0960-9822(00)00061-0 (1994).

200 Feroli, F. *et al.* Analysis of a 17.9 kb region from Saccharomyces cerevisiae chromosome VII reveals the presence of eight open reading frames, including BRF1 (TFIIIB70) and GCN5 genes. *Yeast* **13**, 373-377, doi:10.1002/(SICI)1097-0061(19970330)13:4<373::AID-YEA82>3.0.CO;2-V (1997).

201 Ciriacy, M. Isolation and characterization of yeast mutants defective in intermediary carbon metabolism and in carbon catabolite derepression. *Mol Gen Genet* **154**, 213-220, doi:10.1007/BF00330840 (1977).

202 Bourgeron, T. *et al.* Mutation of a nuclear succinate dehydrogenase gene results in mitochondrial respiratory chain deficiency. *Nat Genet* **11**, 144-149, doi:10.1038/ng1095-144 (1995).

203 Lee, Y. J., Jang, J. W., Kim, K. J. & Maeng, P. J. TCA cycle-independent acetate metabolism via the glyoxylate cycle in Saccharomyces cerevisiae. *Yeast* **28**, 153-166, doi:10.1002/yea.1828 (2011).

204 Parsell, D. A., Kowal, A. S., Singer, M. A. & Lindquist, S. Protein disaggregation mediated by heat-shock protein Hsp104. *Nature* **372**, 475-478, doi:10.1038/372475a0 (1994).

205 Parsell, D. A., Sanchez, Y., Stitzel, J. D. & Lindquist, S. Hsp104 is a highly conserved protein with two essential nucleotide-binding sites. *Nature* **353**, 270-273, doi:10.1038/353270a0 (1991).

206 Sanchez, Y., Taulien, J., Borkovich, K. A. & Lindquist, S. Hsp104 is required for tolerance to many forms of stress. *EMBO J* **11**, 2357-2364 (1992).

207 Gammie, A. E., Stewart, B. G., Scott, C. F. & Rose, M. D. The two forms of karyogamy transcription factor Kar4p are regulated by differential initiation of transcription, translation, and protein turnover. *Mol Cell Biol* **19**, 817-825, doi:10.1128/MCB.19.1.817 (1999).

208 Kurihara, L. J., Stewart, B. G., Gammie, A. E. & Rose, M. D. Kar4p, a karyogamy-specific component of the yeast pheromone response pathway. *Mol Cell Biol* **16**, 3990-4002, doi:10.1128/MCB.16.8.3990 (1996).

209 Tripodi, F., Fraschini, R., Zocchi, M., Reghellin, V. & Coccetti, P. Snf1/AMPK is involved in the mitotic spindle alignment in Saccharomyces cerevisiae. *Sci Rep* **8**, 5853, doi:10.1038/s41598-018-24252-y (2018).

210 Lahav, R., Gammie, A., Tavazoie, S. & Rose, M. D. Role of transcription factor Kar4 in regulating downstream events in the Saccharomyces cerevisiae pheromone response pathway. *Mol Cell Biol* **27**, 818-829, doi:10.1128/MCB.00439-06 (2007).

211 Ohtake, Y. & Wickner, R. B. Yeast virus propagation depends critically on free 60S ribosomal subunit concentration. *Mol Cell Biol* **15**, 2772-2781, doi:10.1128/MCB.15.5.2772 (1995).

212 Nicastro, R. *et al.* Snf1 Phosphorylates Adenylate Cyclase and Negatively Regulates Protein Kinase A-dependent Transcription in Saccharomyces cerevisiae. *J Biol Chem* **290**, 24715-24726, doi:10.1074/jbc.M115.658005 (2015).

213 Hoose, S. A. *et al.* A systematic analysis of cell cycle regulators in yeast reveals that most factors act independently of cell size to control initiation of division. *PLoS Genet* **8**, e1002590, doi:10.1371/journal.pgen.1002590 (2012).

214 Cummins, C. M., Gaber, R. F., Culbertson, M. R., Mann, R. & Fink, G. R. Frameshift suppression in Saccharomyces cerevisiae. III. Isolation and genetic properties of group III suppressors. *Genetics* **95**, 855-879 (1980).

215 Dascher, C., Ossig, R., Gallwitz, D. & Schmitt, H. D. Identification and structure of four yeast genes (SLY) that are able to suppress the functional loss of YPT1, a member of the RAS superfamily. *Mol Cell Biol* **11**, 872-885, doi:10.1128/mcb.11.2.872-885.1991 (1991).

216 Margulis, N. G. *et al.* Analysis of COPII Vesicles Indicates a Role for the Emp47-Ssp120 Complex in Transport of Cell Surface Glycoproteins. *Traffic* **17**, 191-210, doi:10.1111/tra.12356 (2016).

217 Fernandez, G. E. & Payne, G. S. Laa1p, a conserved AP-1 accessory protein important for AP-1 localization in yeast. *Mol Biol Cell* **17**, 3304-3317, doi:10.1091/mbc.e06-02-0096 (2006).

218 Yashiroda, H., Kaida, D., Toh-e, A. & Kikuchi, Y. The PY-motif of Bul1 protein is essential for growth of Saccharomyces cerevisiae under various stress conditions. *Gene* **225**, 39-46, doi:10.1016/s0378-1119(98)00535-6 (1998).

219 Qian, W., Ma, D., Xiao, C., Wang, Z. & Zhang, J. The genomic landscape and evolutionary resolution of antagonistic pleiotropy in yeast. *Cell Rep* **2**, 1399-1410, doi:10.1016/j.celrep.2012.09.017 (2012).

220 Watanabe, M., Watanabe, D., Nogami, S., Morishita, S. & Ohya, Y. Comprehensive and quantitative analysis of yeast deletion mutants defective in apical and isotropic bud growth. *Curr Genet* **55**, 365-380, doi:10.1007/s00294-009-0251-0 (2009).

221 Amerik, A. Y., Li, S. J. & Hochstrasser, M. Analysis of the deubiquitinating enzymes of the yeast Saccharomyces cerevisiae. *Biol Chem* **381**, 981-992, doi:10.1515/BC.2000.121 (2000).

222 Bohm, S. *et al.* The Budding Yeast Ubiquitin Protease Ubp7 Is a Novel Component Involved in S Phase Progression. *J Biol Chem* **291**, 4442-4452, doi:10.1074/jbc.M115.671057 (2016).

223 Enenkel, C., Blobel, G. & Rexach, M. Identification of a yeast karyopherin heterodimer that targets import substrate to mammalian nuclear pore complexes. *J Biol Chem* **270**, 16499-16502, doi:10.1074/jbc.270.28.16499 (1995).

224 MacKinnon, M. A. *et al.* The Kap60-Kap95 karyopherin complex directly regulates phosphatidylcholine synthesis. *J Biol Chem* **284**, 7376-7384, doi:10.1074/jbc.M809117200 (2009).

225 Kreike, J., Schulze, M., Pillar, T., Korte, A. & Rodel, G. Cloning of a nuclear gene MRS1 involved in the excision of a single group I intron (bI3) from the mitochondrial COB transcript in S. cerevisiae. *Curr Genet* **11**, 185-191, doi:10.1007/BF00420605 (1986).

226 Benito-Moreno, R. M., Miaczynska, M., Bauer, B. E., Schweyen, R. J. & Ragnini, A. Mrs6p, the yeast homologue of the mammalian choroideraemia protein: immunological evidence for its function as the Ypt1p Rab escort protein. *Curr Genet* **27**, 23-25, doi:10.1007/BF00326574 (1994).

227 Singh, J. & Tyers, M. A Rab escort protein integrates the secretion system with TOR signaling and ribosome biogenesis. *Genes Dev* **23**, 1944-1958, doi:10.1101/gad.1804409 (2009).

228 Jamalzadeh, S., Pujari, A. N. & Cullen, P. J. A Rab escort protein regulates the MAPK pathway that controls filamentous growth in yeast. *Sci Rep* **10**, 22184, doi:10.1038/s41598-020-78470-4 (2020).
